# Development of a UVGI System and Evaluation of Germicidal Potential Against Biofilm-Forming Bacteria and Fungi Under Controlled Laboratory Conditions

**DOI:** 10.64898/2026.03.31.715580

**Authors:** Bindu Sadanandan, Shyam Sunder, Vaniyamparambath Vijayalakshmi, Priya Ashrit, Kavyasree Marabanahalli Yogendraiah, Kalidas Shetty

**Affiliations:** M S Ramaiah Institute of Technology, Department of Biotechnology, Bangalore 560054, Karnataka, India; Bharat Electronics Limited, Product Development & Innovation Center, Bangalore 560013, Karnataka, India; North Dakota State University, Department of Microbiological Sciences, Fargo, ND58105, USA

**Keywords:** UVGI, UVC, Sterilization, Biofilm, Environmental Sanitation

## Abstract

A compact, in-house developed ultraviolet germicidal irradiation (UVGI) system adaptable to static, mobile, or robotic platforms was developed for the effective sterilization of bacteria and fungi using a wireless mode of operation. Under controlled laboratory conditions, its efficacy was evaluated against three representative biofilm-forming pathogens: *Bacillus subtilis* (Gram-positive, spore-forming, motile bacterium), *Escherichia coli* K12 (Gram-negative, non-spore-forming, non-motile bacterium), and *Candida albicans* M-207 (multi-drug-resistant, clinical yeast isolate). Microbial viability following UVGI exposure was assessed using colony-forming unit (CFU) and MTT assays, and morphological alterations were characterized by scanning electron microscopy (SEM). Cultures were exposed to UV-C radiation at distances of 1–5 m for 15–90 min. CFU assay demonstrated 100% kill of all tested organisms at 1 m and 15 min, corresponding to doses of 600.3, 576 & 697.5 mJ/cm². Although MTT assays indicated residual metabolic activity under the same conditions, CFU results confirmed that surviving cells were unable to proliferate, highlighting the robustness of UV treatment for long-term inactivation. SEM confirmed distinct morphological alterations such as complete destruction of extracellular matrix & reduction in number of cells indicating cell death with increase in UV dose as compared to controls. A dose & time-dependent inactivation of biofilm-forming bacteria & fungi was observed on exposure to UVGI. Therefore, this pilot study validates the effectiveness of the newly developed UVGI surface sterilizer against biofilm-forming bacterial and fungal pathogens. Overall, the system demonstrates proof-of-concept efficacy under laboratory conditions and holds strong potential for future development and validation in hospitals and other contaminated public spaces.

**Graphical Abstract:** 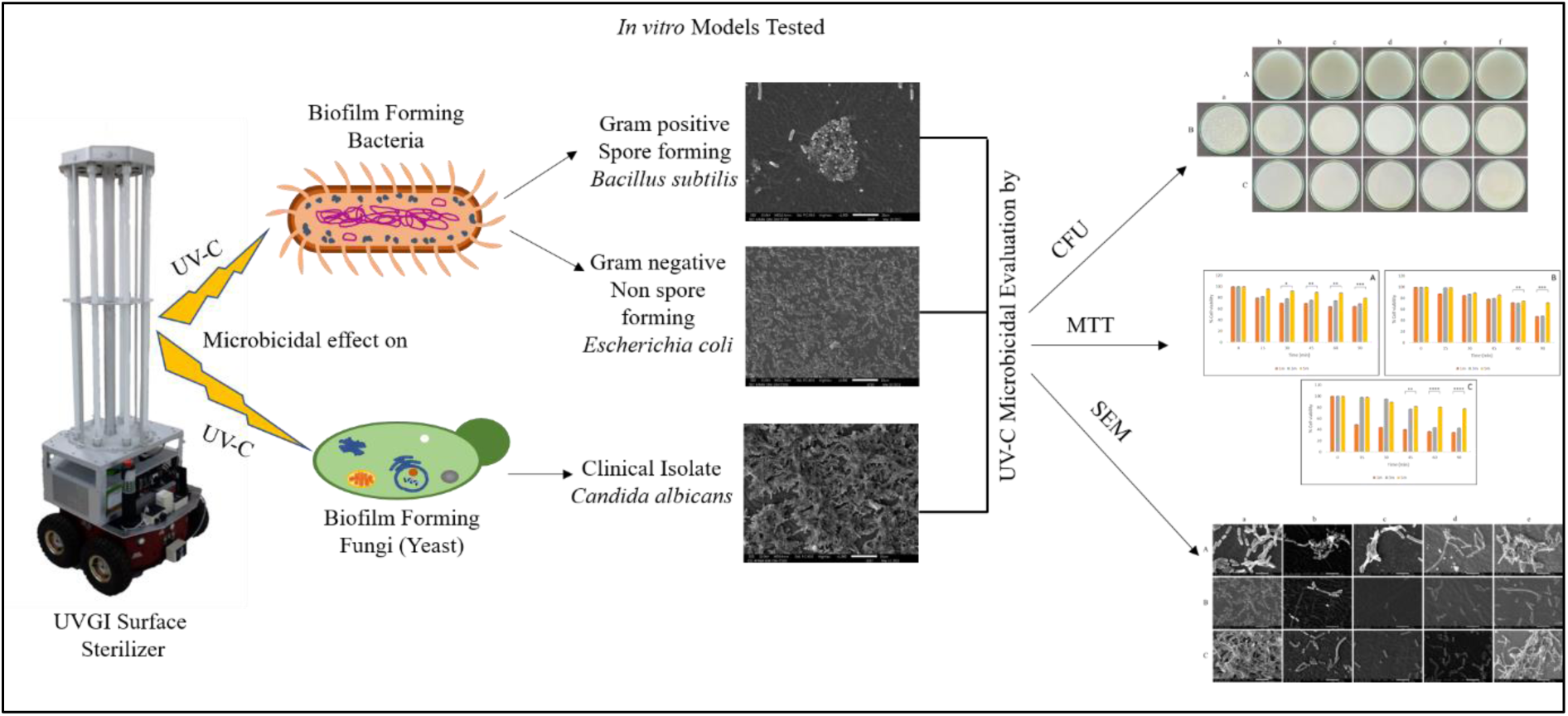

**IMPORTANCE:** Microorganisms that form biofilms on surfaces are difficult to eliminate and contribute to the spread of infections in healthcare and indoor environments. There is a need for practical, easy-to-use disinfection technologies that can effectively reduce such contamination. In this study, we developed a compact, in-house, wireless UV-C disinfection system designed for flexible operation across different surface types. The system was evaluated under controlled laboratory conditions using representative biofilm-forming bacterial and fungal pathogens. Our findings show that the system can effectively reduce microbial contamination, demonstrating proof-of-concept efficacy. This work highlights the potential of accessible, non-chemical UV-based technologies and supports their further validation for applications in real-world disinfection settings.

## INTRODUCTION

Microorganisms have been studied in diverse public areas such as telephone booths, toilets, schools, hospitals, and common transportation points (1,2). The highest level of bacterial and fungal contamination was found in hospitals, followed by ambulances, indoor environments (kitchen, toilets, schools, restaurants, malls, etc.), and heating/air conditioning systems. In the hospital environment, microorganisms, especially nosocomial pathogens, cause hospital-acquired infections such as Central line-associated blood stream infections (CLABSI), Catheter-associated urinary tract infection (CAUTI), Ventilator-associated pneumonia (VAP), Surgical site infection (SSI), which are of global concern (3). The most common nosocomial pathogen is *Escherichia coli*, which is a common cause of tract infections. Other pathogenic species, such as *Staphylococcus, Klebsiella, Enterobacter, Streptococcus, and Candida,* also cause hospital-acquired infections (4).

Several of these pathogenic bacteria and fungi form structures known as biofilms, which are complex amalgamations of cells that are embedded in a matrix of exopolysaccharides (5), providing them with advantages for survival and infection. It has been over 70 years since the first report on biofilms (6), and biofilms remain elusive owing to their complex structure. These biofilms attach to different surfaces, provide advantages for disseminating infections, and are challenging to eradicate. Approximately 90% of microorganisms live in a biofilm, and only 10% are in their planktonic form (7). If the biofilms dry, then removing them from the surface requires multiple wipes, and the microbial cultures from these biofilms can spread to the next surface (8). Disinfection and sterilization of surfaces and materials such as silicone elastomers, which are used as silicone fillers/gels, and patches for wound healing are essential, especially in hospitals where the dissemination of infection is higher due to the ease of spreading of infections (9).

Healthcare-associated infections remain a significant challenge due to the persistence of microorganisms on environmental surfaces, particularly in the form of biofilms, which exhibit increased resistance to conventional disinfection methods. The COVID-19 pandemic has further emphasized the importance of effective surface decontamination strategies to limit the transmission of infectious agents in both healthcare and public settings (10). Ultraviolet (UV) radiation or UV Germicidal Irradiation (UVGI) is used as an effective physical source for sterilization (11). UV irradiation has been shown to slow biofilm formation (7).

UV radiation is categorized as UV-A (320-400 nm), UV-B (280-320 nm), and UV-C (200–280 nm); UV-C is considered to have germicidal properties (8, 12–14). UVGI is a proven physical technique for disinfecting surfaces that are contaminated with microbes. When irradiated with a UV-C source, inactivation is due to photo biochemical reactions that take place within the cell. The effectiveness of UV-C is attributed to its ability to damage the Deoxyribonucleic Acid (DNA) by altering nucleotide base pairing. Protein molecules, along with DNA and Ribonucleic acid (RNA), absorb UV photons between 200-300 nm, with peak absorption at 260 nm, thus destroying the cells (15, 16).

The development of a UVGI system is complex, and the challenge lies in the transportation of the system and the manpower involved in the operation of the equipment. Gram-positive and gram-negative bacteria behave differently and have different infectious and toxigenic effects on humans due to their different cellular mechanisms approach (17). The UV-C radiation on these bacteria will be different, as the cell walls of Gram-positive and Gram-negative bacteria are different (18). To overcome these bacterial infections and clinical yeast isolates, an easy-to-use wireless-based UVGI can be targeted for application globally in the rapid surface sterilization of hospitals, culture rooms, restaurants, or other contaminated closed spaces, and can contribute to the rapid management of these associated infectious agents, reducing their rapid transmission.

The motivation for this work stems from several critical research gaps in the current landscape of ultraviolet germicidal irradiation (UVGI) technology. Existing commercial UVGI systems are often expensive, rely on proprietary technology, and are typically fixed installations, making them unsuitable for deployment in resource-limited or rural healthcare environments. Additionally, their bulky design and the need for significant manpower to operate and relocate them limit their portability and flexibility, particularly in dynamic settings such as hospitals, clinics, and public spaces. Operational complexity further compounds these challenges, as many systems require trained technicians and involve intricate protocols, which hinder rapid deployment during infectious disease outbreaks. Moreover, while UVGI has demonstrated promising disinfection efficacy, its effectiveness against resilient biofilm-forming bacteria and fungi, especially when tested on clinically relevant substrates such as petri plates, 96-well microtiter plates, and silicone elastomers across different experimental models, remains inadequately explored.

To address these gaps, the present study involves the design and development of a compact, in-house wireless UVGI system integrating hardware and control software for remote operation via a mobile device or computer over a WiFi network. The antimicrobial efficacy of the system was evaluated using representative biofilm-forming microorganisms, including *Bacillus subtilis* (Gram-positive, spore-forming), *Escherichia coli* K12 (Gram-negative), and a multidrug-resistant clinical isolate of *Candida albicans* M-207. The study systematically investigates the effect of UV-C exposure at varying distances and time intervals using colony-forming unit (CFU) enumeration, MTT assay, and scanning electron microscopy (SEM) analysis. The findings aim to provide a comprehensive assessment of UVGI efficacy against biofilm-associated microorganisms and to evaluate the practical applicability of a wireless UVGI platform for surface disinfection.

## METHODOLOGY

### Development of the UVGI surface sterilizer system

Commercially available UVGI equipment has the disadvantage of having a complex structure, being fixed and suitable only for targeted and specific room sterilization, and having proprietary technology; therefore, servicing is also expensive, especially affecting rural health centers around the world. The UVGI system developed in this study is a simple assembly setup that is convenient, portable, and can be operated in wireless mode. The UVGI system was developed with two basic components: hardware and software. The hardware component consists of UV tubes, chargers, batteries, inverters, Node Microcontroller Unit (MCU) hardware, an aluminium structure to hold everything in place, and rubber gaskets.

UV dose values are calculated as: Formulae used: Dose = Intensity × Exposure Time Example : *E. coli* UV dose values are calculated as follows:

- At 1 m, 0.64 mW/cm², 15 min → 576 mJ/cm²
- At 3 m, 0.177 mW/cm², 30 min → 318.6 mJ/cm²
- At 5 m, 0.032 mW/cm², 90 min → 172.8 mJ/cm²

The Dosimeter is factory calibrated

A dosimeter (Model: RMD Radiometer -814400C, Make: Opsytec, Dr. Gröbel Gmbh, Germany) was used as an external system to measure the intensity of UV radiation. The software is an in-built application in the Node MCU System on Module (SoM).

- Lamp type and peak wavelength: Philips UV-C germicidal tubes, 36 W, with a peak emission at 253.7 nm.
- Irradiance uniformity: The irradiance was measured at sample positions (1 m, 3 m, 5 m) using a calibrated dosimeter, and uniformity across the exposure plane was verified by measuring at multiple points, showing <10% variation in irradiance.
- Lamp lifetime: The manufacturer specifies an operational lifetime of ∼9,000 hours at 80% rated output.
- Emission characteristics: The tubes have a sharp emission peak at 253.7 nm, with negligible emission below 240 nm, making them ozone-free.

The UVGI system was developed considering the requirements of reduced manpower, portability, and simple applications. The UVGI was structurally assembled using locally sourced materials that can be easily mass-produced if needed. Any person with a basic knowledge of electronics can assemble it together; hence, the labour hours and cost for assembling is low. This device can be used for sterilization and basic training.

### Microbial strains

Three different biofilm-forming model microbial systems were selected for this study. Gram-positive spore-forming bacteria, *Bacillus subtilis* MTCC 441 (ATCC-6633), and a gram-negative non-spore-forming bacterium, *Escherichia coli* K12 (MTCC–1302), were procured from IMTECH, Chandigarh, India. The Yeast: *Candida albicans* M-207, a fluconazole and caspofungin multi-drug resistant clinical isolate from the umbilical vein catheter of a female baby with invasive *Candidiasis,* was provided by the Department of Microbiology, M S Ramaiah Medical College and Teaching Hospital, Bengaluru, India. No human subjects were directly involved in the study; hence, ethical clearance/consent was not required.

We previously used *C. albicans* M-207 clinical isolate as a model system in subsequent studies. The growth conditions of *Candida* species, including *C. albicans* M-207, have been optimized using design of experiments (Response Surface Methodology (RSM)) (19). The growth media for *C. albicans* M-207 and *E. coli* ATCC 39936 in a polymicrobial association have also been optimized using RSM (20). *C. albicans* M-207 was found to be Multidrug resistant (MDR) (Fluconazole and Caspofungin); hence, studies on the control of *C. albicans* M-207 (21) and its polymicrobial interaction with *E. coli* (22) have been conducted with aqueous spice extracts (Garlic, Indian gooseberry, and clove).

### Growth conditions

*B. subtilis* and *E. coli* K12 were streaked on Nutrient Agar and incubated at 37°C for 24 h, whereas Yeast Extract Peptone Dextrose (YEPD) agar was used to maintain *C. albicans* M-207 at 32°C for 24 h. Colonies were picked from these plates and used for further studies. The culture plates were sub-cultured every 15 days. Glycerol stocks (15% v/v) of the cultures were maintained at -20°C for short-term storage, and the mother cultures were maintained at -86°C.

### Substrate Material

In two of the test models, silicone elastomer discs were used as substrates to grow the bacteria and yeast cultures, as this material is widely used in hospitals in medical devices such as catheters, liners for prostheses, valves, tubing’s, and long-term implants owing to its biocompatibility. Studying the effect of UV-C on cells grown on silicone elastomer materials will be helpful in controlling the formation of biofilms on medical devices made of this material.

### Agar Spot Assay

A loopful of each of the cultures of *B. subtilis, E. coli* K12, and *C. albicans* M207 in separate Petri plates was inoculated at the centre of TSA media in separate petri plates. The plates were incubated at 37°C for 16 h.

### Preparation of Inoculum

The pre-inoculum was prepared from subcultured plates showing biofilm growth & a loopful of the culture was inoculated in the respective broth media (nutrient broth for *B. subtilis* and *E. coli* K12 and YEPD for *C. albicans* M-207) in a test tube and incubated for 16 h in an incubator shaker at 100 rpm at 37^0^C.

Though preinoculum is prepared in liquid media. we have maintained the shaker speed of 100 rpm to control the and promote biofilm growth. Biofilm formation has been previously optimized. While shear forces in liquid culture limit surface attachment, microcolonies and suspended aggregates (non-attached biofilm fragments) are still present. In static or low-shear liquid cultures, biofilms form on the bottom and sides of wells or tubes, rather than freely suspended in the medium. Thus, preinoculum retains biofilm characteristics and, when added onto silicone elastomer discs, they develop into mature surface biofilms. Therefore, the UV-C disinfection experiments were performed on established biofilms, consistent with the study objectives and relevant to real-world biofilm-associated microbial contamination.

### Petri plate model to assess viable count of UVGI irradiated cultures - Colony Forming Unit

The microbial inoculum for the UVGI experiments was standardized to 1 × 10⁶ CFU/mL for all organisms to ensure consistent initial cell loading and reproducible formation of surface-attached communities under experimental conditions. Serial dilutions of actively growing cells were prepared using a slightly modified protocol (23). A 100 µL volume of 10^−5^ dilution of the bacterial cultures and 10^−2^ dilution of the yeast culture were added to the NA and YEPD plates, respectively, and spread evenly on the agar surface using a sterile spreader. The different dilutions for bacterial and yeast cultures were used to ensure well-defined countable colonies that are statistically significant (30–300) after plating for CFU determination. Petri plates were placed inside the laminar airflow chamber and exposed to different intensities of UV-C emitted by the UVGI sterilizer at distances of 1, 3, and 5 m with a UV dose of 600.3, 576 & 697.5 mJ/cm². Petriplates were placed perpendicular to the tubelight mounted vertically on the robot and parallel to the Laminar air flow (LAF) bench. The intensities were measured at all distances using a UV dosimeter. The plates were exposed to UV-C for 15, 30, 45, 60, or 90 min at each distance. The control sets were maintained in parallel. Control sets are plates with microbial culture without UV exposure. The plates are inoculated with the culture and incubated in the incubator, just like the ones after the UV exposure. All experiments were performed in triplicates. After each exposure, the plates were sealed and incubated at 37°C for 18 h. After incubation, the colonies were counted and Colony Forming Unit (CFU) CFUmL^−1^ was calculated using the formula (Eq. 1). Percent kill/percentage reduction was calculated by the formula (Eq. 2)

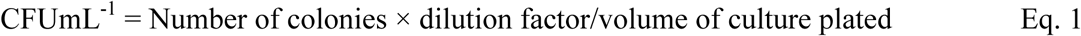

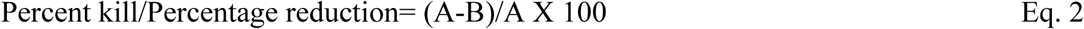

where A is the number of viable bacteria before UV irradiation

B is the number of viable bacteria after UV irradiation

### Silicone elastomer disc in 96-well microtiter plate model to assess viability of the cultures exposed to UVGI – MTT (3-(4,5-Dimethylthiazol-2-yl)-2,5-Diphenyltetrazolium Bromide) Assay

Sterile 5 mm diameter silicone elastomer discs were placed in separate wells of 96-well flat-bottom polystyrene microtiter plates. The inoculum (100 µL) was added to each well of the microtiter plate and incubated at 37°C for 90 min. After 90 min, 1 × phosphate buffered saline (PBS) was added, and the plates were exposed to UV-C for 15, 30, 45, 60, and 90 min at 1, 3, and 5 m distances. Control plates of microbial culture without UV-C irradiation were maintained in parallel. During UV exposure, samples were held in phosphate-buffered saline (PBS), which provides an optically clear, isotonic medium with minimal UV absorption. This ensured the delivery of a defined UV dose to the microbes while minimizing attenuation by organic material. After exposure, the PBS was gently removed to remove non-adherent planktonic cells, and 100 µL of fresh growth medium was added to recover and resuspend the remaining attached biofilm cells. The plates were then incubated for 24h at 37 °C & then, The MTT assay was performed (24). After the incubation period, the medium was discarded and the wells were washed with PBS. A 50 µL volume of 5 mgmL^−1^ MTT solution was added to each well and incubated for 3 h. After incubation, 100 µL of acidified isopropanol was added to each well to dissolve formazan crystals and incubated for 10-20 min. The wells were mixed thoroughly, 100 µL of the mixture was transferred to a fresh well, and the absorbance was read at 540 nm using a Synergy HT microtiter plate reader.

### Silicone elastomer discs in petri plate model for morphological analysis of the test organisms on irradiation - Scanning Electron Microscopy

Biofilms were established on sterile silicone elastomer discs using standardized inoculum. Cultures were revived from stock, subcultured on NA or YPD plates, and loopful growth was transferred into 5 mL NB or YPD broth (pre-inoculum) and incubated overnight (37 °C for *E. coli & B. subtilis* & 32 °C for *C. albicans*). The pre-inoculum was diluted 1:60, adjusted to 1 × 10⁶ cells/mL, and added onto the discs under identical incubation conditions. After growth, non-adherent cells were gently removed by PBS washing, leaving only surface-attached biofilms. This procedure ensured reproducible inoculum density and minimized variability in baseline biofilm formation across replicates.

Serially diluted cultures of bacteria and yeast were prepared in nutrient broth and YEPD broth, respectively, and poured into petri plates containing 15 mm diameter silicone elastomer discs. The petri plates with silicone discs were exposed to UV-C for different durations and distances. The Petriplates were then incubated overnight. The control set was maintained in parallel. After incubation, the discs were washed with PBS and fixed using 4% glutaraldehyde for 1 h. Subsequently, the discs were washed with PBS before sequential dehydration in an ethanol series: 70% for 10 min, 95% for 10 min, and 100% for 20 min (21). The discs were air-dried completely, coated with gold under vacuum, and visualized at 5000x & 1900x, 15 kV using a JEOL IT 300 Scanning Electron Microscope at AFMM, Indian Institute of Science, Bengaluru.

### Statistical Analysis

Three independent tests were performed to ensure the reliability and reproducibility of the data. All experimental data are expressed as the mean ± standard deviation. Sextuplicates were maintained for each of the experiments involving microtiter plates, and triplicates were maintained for the other experiments. For cell viability, the data are presented as a percentage of the control, and the significance with respect to the control is presented. For percentage reduction, the significance of 3m and 5m with respect to 1m was determined. Two-way analysis of variance (ANOVA) and Tukey’s multiple comparison tests were performed using GraphPad Prism 9.

Linear regression analysis and the calculation of the coefficient of determination (R²) are commonly used statistical tools to evaluate the strength and linearity of a relationship between two variables. However, in microbiological experiments, especially those involving microbial growth, biofilm formation, disinfection kinetics, biological variability, non-linear growth dynamics, and environmental influences often make it difficult to obtain a high R² value (close to 1). Microbial responses are inherently complex and influenced by multiple factors that do not always follow a simple linear trend, so linear regression is not typically suitable for interpreting experimental outcomes in such systems.

## RESULTS

### Device Development

#### Hardware

The UVGI disinfection system consisted of eight UV tubes (Philips, India), each of 36 W, that were energized with their own ballast. The power supply from the inverter via the AC relay was supplied to each ballast, which is an energizing element that powers the UV tube. The ballast requires a 40 W input to provide 36 W to the UV tube and the remaining for its own power. Hence, a total of 400 W of input power was required, considering the inverter efficacy to be 75-80% and other losses. It was essential to provide AC power as a backup for the system, which was achieved by combining an inverter, charger, and 24 V battery system. The battery was a 50 Ah battery from Micronix Ltd., a Bangalore-based micro-, small-, and medium-sized enterprise (MSME). The inverter (Meanwell) was a 24 V DC to 240 V auto-grade AC square-wave converter with an inbuilt short circuit and under/over-voltage protection. The battery was 24 V DC, 50 Ah, and a Li-Ion type battery with a charger. Together, this provides a 3 h power backup. A block diagram of the UVGI system is depicted in Figure 1 (a), and the overall system specifications are listed in Supplementary Table 1.

**Figure 1.**
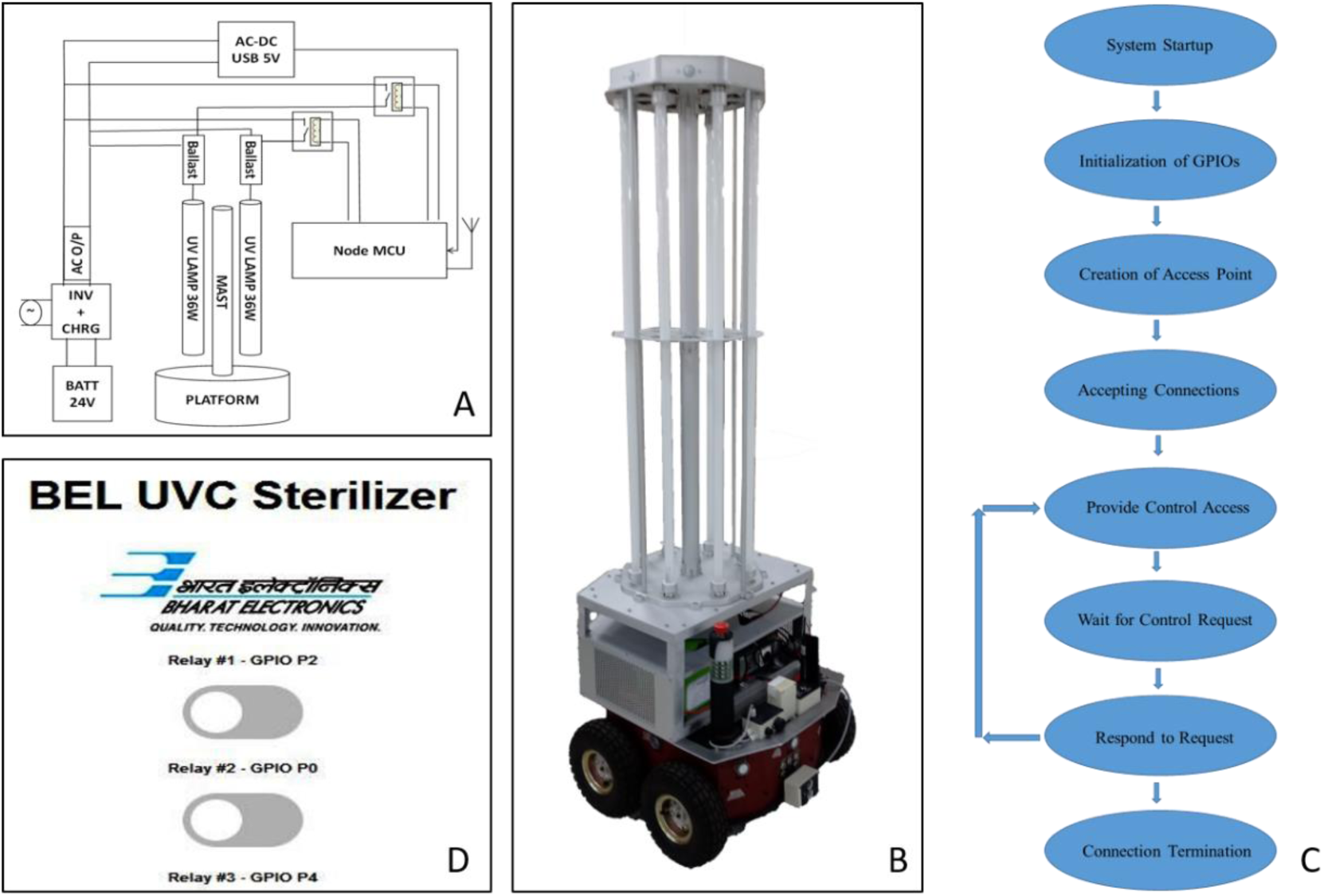
(A) Block Diagram of UVGI Disinfection System, (B) Mechanical Setup of UVGI holding UV tubes arranged on a vehicle, (C) Systematic flow of the software, (D) Control Page Hosted on Node MCU for Control of the UVGI system

The eight UV tubes were supported with an aluminium (Alloy 6061-T6) structure (MAST) designed for easy machining without compromising the required strength. The aluminium alloy was further coated with an 18 micron-thick and clear anodic coating to prevent corrosion and reduce damage due to rough handling. The UV tubes were secured in the middle with a centre plate and cylindrical tube holders at their ends. Rubber gaskets with a shore hardness of 65±7 A were fixed between the metal and tube interface to avoid breakage owing to stress. An aluminium extruded hollow rod with 2-inch outer diameter and 3 mm was fixed at the centre to provide stiffness to the structure. Four rods with a half-inch diameter were placed around the periphery to improve the stability of the UVGI system. The dosage of UV-C on the surface was measured using a sensor interfaced with a digital interface meter known as a dosimeter. The complete setup of UVGI is shown in Figure 1 (b).

In the current study, the UVGI system employed 36 W Philips UV-C tubes with a peak emission at 253.7 nm, which are standard low-pressure mercury discharge lamps. These lamps are widely used in germicidal applications due to their strong emission near 254 nm, the wavelength of maximum DNA absorption. The system did not include an inbuilt ozone generator, and ozone concentration was not measured during the experiments. However, the Philips UV-C tubes used are commercially classified as “ozone-free,” as their emission spectrum does not include significant radiation below 240 nm, which is required for ozone generation. The emission spectrum provided by the manufacturer indicates a sharp peak at ∼253.7 nm with negligible sideband emissions.

#### Software

The Input/output of each relay was connected to Node MCU hardware, which is an SoM based on Espressif System Platform 32 (ESP32), an Advanced RISC Machine (ARM)-based System on Chip (SoC) having inbuilt Wireless Fidelity Media Access Control (WiFi MAC). The MCU SoM node hosts the webpage through which the user can control the system. An AC-DC adapter with a 5 V output was used to power the SoM module. The Node MCU SoM module had the necessary software to control the system via a web server; therefore, it does not require any secondary software.

When the system was switched on, an access point was created by the hardware that could be connected via WiFi, either on the phone or laptop, to the IP address of the device and copied onto a browser. The browser is then connected to a webpage hosted on the SoM module. Using this web browser, the user can control the device wirelessly, without any direct contact with the device. This protects the user from UV radiation and ensures device safety.

The flow of the software is depicted in Figure 1 (c), which shows the general procedure followed when the system is at the stage of accepting control and providing the connection. Fig. 1 (d) shows a snapshot of the web browser.

#### Operation procedure

- Plug the red wire into the red connector (positive) of the battery and the black wire into the black connector (negative). Never exchanged positions.
- Switch on the battery.
- Connect the UVGI system to the power source and switch on.
- The WiFi network is opened on the mobile/laptop and connected to the IP address of the system.
- Copy the IP address of the network and paste it in the browser.
- The browser then opens the control page to switch the UV tubes on/off using the relays.
- The experiments were performed, the relays were manually switched off, and the system and battery switch were switched off.
- Remove one of the connectors, either positive/negative

#### Safety Paradigm

Despite its formidable disinfection power, UV-C radiation carries inherent risks to human health, necessitating stringent safety protocols because this system is efficient in UV-mediated breaking of the DNA/RNA of an organism. With the primary concern of the potential for skin and ocular damage upon direct exposure, meticulous attention must be paid to safety measures. The system should be handled by trained professionals, and safety gear such as polypropylene face masks, polyester suits, and gloves, which exhibit very low transmissivity properties, should be worn during usage.

#### Safety Protocol for Operating UVC Room Sterilizers

Operation of UVC sterilizers requires strict adherence to safety protocols to protect personnel from harmful exposure to UVC radiation (200–280 nm)

Before the operation, the designated area must be cleared of all occupants. Entry should be restricted with visible warning signs indicating UVC sterilization in progress. Operators must wear personal protective equipment (PPE), including UV-blocking goggles with optical density >5, nitrile gloves, face masks, and full-sleeved UV-resistant garments. A UV protective kit should always be available at the worksite.

During operation, personnel must avoid direct line of sight with active lamps. Remote or wireless control systems should be used to power the unit on and off, minimizing manual intervention. For real-world UVGI applications, strict adherence to safety protocols and PPE use is essential to prevent accidental UV-C exposure, which can damage skin and eyes.

#### Safety Measures Incorporating Specific Personal Protective equipment (PPE) and Goggles Specification

- PPE Material Selection: When selecting personal protective equipment (PPE), prioritizing materials resistant to UV-C radiation is imperative. Typically made of UV-resistant, tightly woven fabrics such as polyester-cotton blends or specialized UV-blocking materials. Garments composed of UV-resistant fabrics offer enhanced protection against skin damage.
- Gloves: Nitrile or neoprene gloves are preferred, as they resist UV degradation better than latex and protect hands during handling of UV equipment or contaminated materials.
- Goggle Specification: UV-resistant goggles or face shields/helmets made from polycarbonate or acrylic materials that block UV-C wavelengths (<280 nm). These prevent photokeratitis (“welder’s eye”) and long-term retinal damage. Utilization of UV-blocking goggles featuring lenses engineered to effectively filter UV-C radiation is paramount. Goggles with a high Optical Density (OD) rating, preferably surpassing OD 5, ensure robust ocular protection by attenuating the UV-C wavelengths. This mitigates the risk of ocular injury during disinfection procedures.
- Footwear: Closed shoes with UV-resistant coatings or standard laboratory safety shoes, which protect against accidental UV reflection off floors.
- Augmented Skin Protection: To fortify skin defences against UV-C exposure, the application of barrier creams or lotions must be considered. Formulations designed to provide comprehensive protection against chemical exposure or contact dermatitis enhance skin resilience and minimize potential irritant effects.
- Exposure Mitigation: Vigilantly mitigate direct exposure risks by refraining from gazing directly at activated UV-C tube lights and ensuring the absence of personnel within the vicinity during disinfection.
- Ventilation vigilance: Implementing robust ventilation mechanisms within the disinfection area to mitigate the accumulation of ozone, a byproduct of UV-C irradiation, is renowned for its deleterious health effects when inhaled at elevated concentrations.

#### Safety Protocols during Accidental exposure

- Operate UVGI systems only in enclosed or controlled environments with interlock mechanisms and warning indicators.
- Workers must never be directly exposed to UV-C lamps while operational.
- In case of accidental exposure:
- Eyes: Immediately close eyes and move away from the source. Seek medical attention if irritation or pain persists.
- Skin: Cover affected area, avoid further UV exposure, and apply soothing creams (e.g., aloe vera); medical consultation may be required.
- Regular UV irradiance monitoring and equipment maintenance ensure safe and consistent operation.

These measures ensure safe handling of UVGI systems while maintaining their efficacy for microbial control.

### Microbial Validation of UVGI Surface Sterilization

#### Agar Spot Assay

The biofilm-forming ability of the test organisms *B. subtilis, E. coli,* and *C. albicans* M-207 was confirmed by agar spot assay. The agar spot assay was used to assess surface-associated growth and radial expansion from a central inoculation point. This method provides a preliminary indication of the ability of the microorganisms to proliferate and spread across a surface, which is often associated with biofilm-forming potential. After 16 h of incubation, the cultures grew and spread completely, forming a lawn on the petri plate by point inoculation using an inoculation loop, as depicted in Figure 2. This confirmed the ability of the cultures to form a high biofilm.

**Figure 2.**
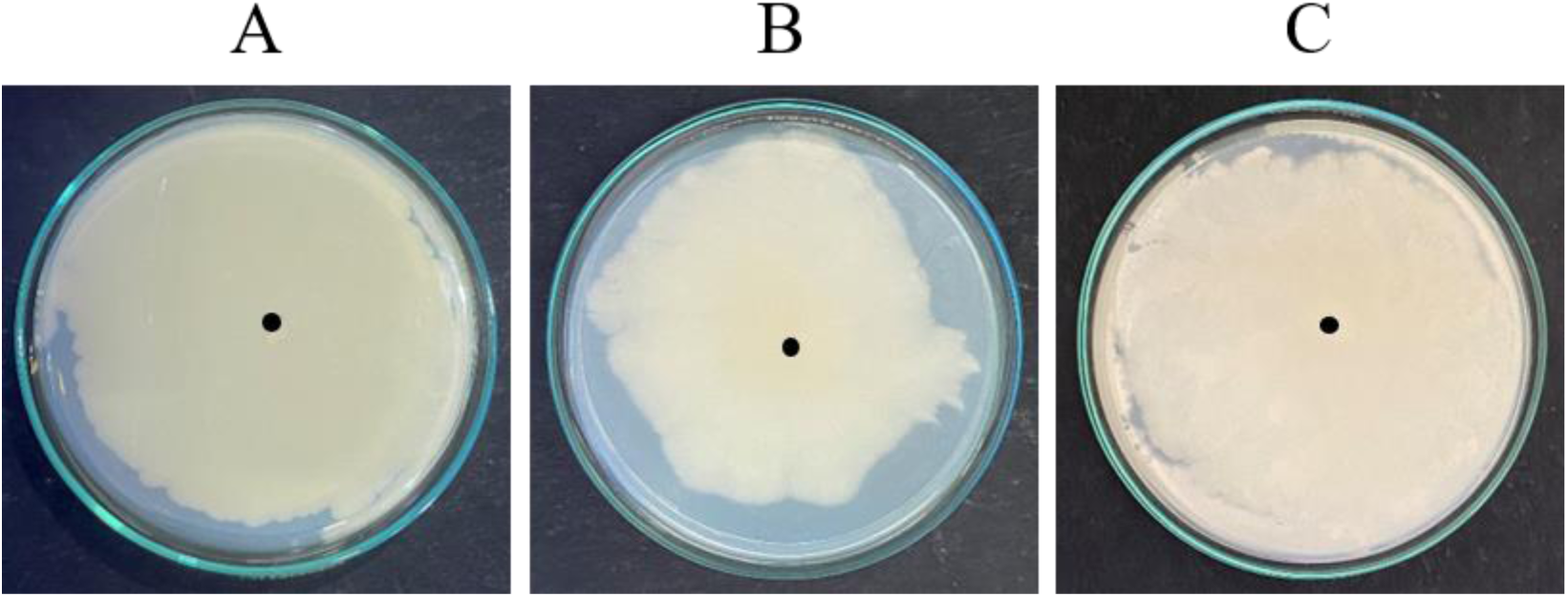
Agar Spot Assay at 16 h. (A) *B. subtilis,* (B) *E. coli* K12, and (C) *C. albicans* M-207. The black dot is the spot of inoculation. The cultures grew and spread completely, forming a lawn on the petri plate. This confirmed the ability of the cultures to form a high biofilm.

#### Colony Forming Unit

The bacterial species *B. subtilis* and *E. coli* K12 were grown in Nutrient Agar medium and the species *C. albicans* M-207 was grown in YEPD medium and irradiated at different time intervals and distances to understand the effect of UV-C radiation emitted by UVGI in inhibiting the growth of these cultures.

The UVGI system effectively killed 100% of the *B. subtilis*, *E. coli* K12, and *C. albicans* M-207 cells at an exposure distance of 1 m and time of 15 min with an UV dose of 600.3, 576 & 697.5 mJ/cm² . At 3m an UV dose of 320.6, 318.6, 318.6 mJ/cm² and an exposure time of 30 min, 99.60% kill was observed for *B. subtilis* & *E.coli* & for *C. albicans* M207 it is 99.89%. Even, at 5 m with an UV dose of 147.6, 477.9 & 637.2 mJ/cm², the percentage kill of cells at different time intervals for *B. subtilis*, *E. coli* K12 & *C. albicans* M-207 was between 99.20-100%. But only at 5m for *C. albicans* M-207, with an UV dose of 469.8 mJ/cm² and an exposure time of 90, showed 65.91 % killing of cells as compared to the control (Figure 3-6). In *C. albicans* M-207, we observed a very interesting phenomenon of the formation of crescent-shaped aggregates of colonies at all time durations at a distance of 5 m. The UV dose and the number of cells (log CFU mL^−1^) are tabulated in Table 1 for 1, 3, and 5 m for *B. subtilis, E. coli* K12, and *C. albicans* M-207, respectively. The effect of UV irradiation on the cell viability & inactivation of biofilm-forming bacteria & fungi was evaluated using colony-forming unit (CFU) counts as shown in Table 2. UVGI treatment resulted in a significant reduction in CFU across (% kill) at different exposure distances and time intervals. The reduction was more pronounced at shorter distances and longer exposure times, with statistically significant differences observed compared to 1 m distance (*p ≤ 0.05–0.01*), as determined by two-way ANOVA followed by Tukey’s post-hoc test.

**Figure 3.**
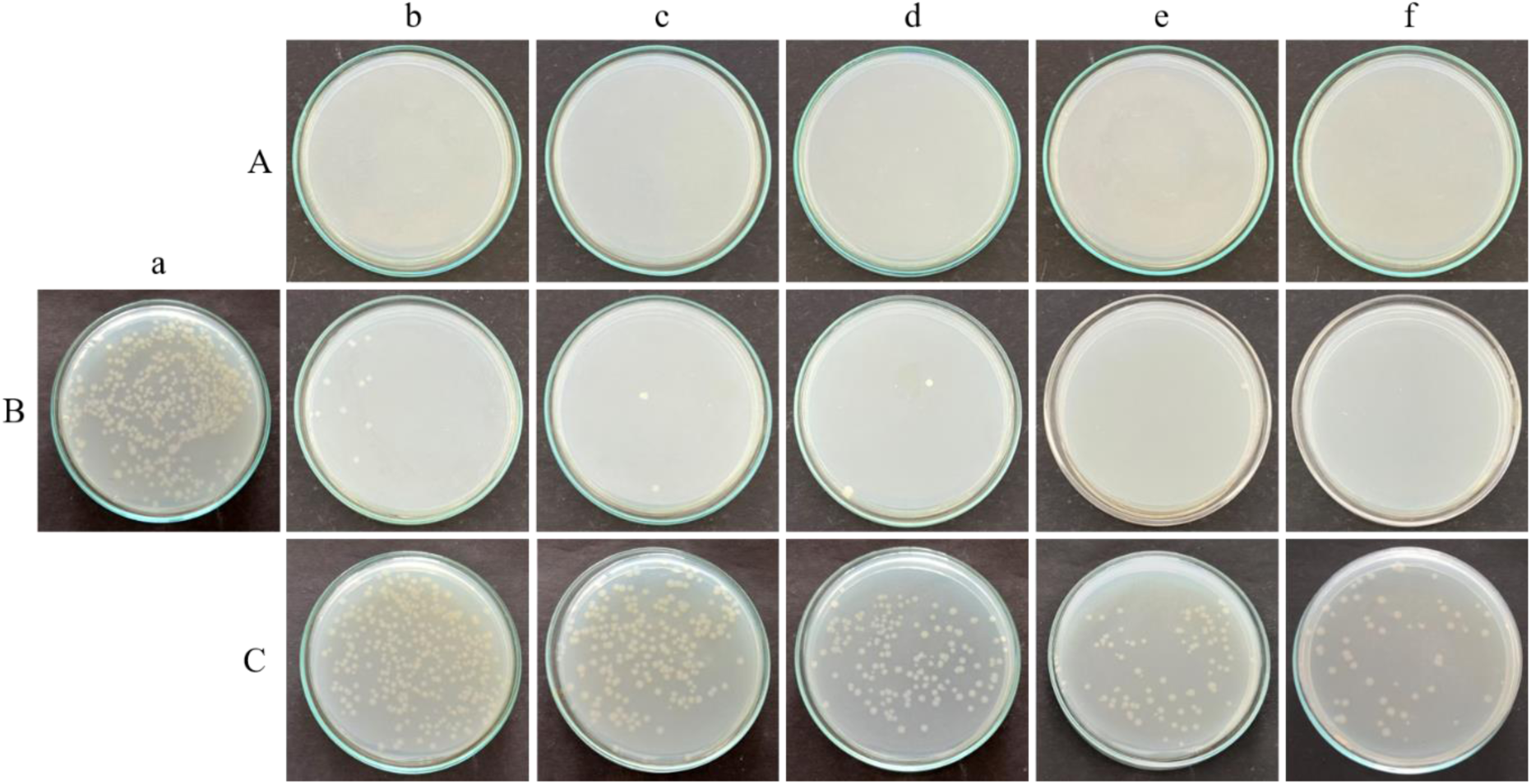
Effect of UV irradiation on E. coli K12 assessed by CFU assay on nutrient agar plates. Bacterial cultures were exposed to UV light at three different exposure distances: (A) 1 m, (B) 3 m, and (C) 5 m. For each distance, plates show (a) untreated control (no UV exposure) and samples exposed for (b) 15 min, (c) 30 min, (d) 45 min, (e) 60 min, and (f) 90 min. A progressive decrease in CFU density with increasing exposure time and proximity to the UV source demonstrates the dose- and distance-dependent bactericidal effect of UV irradiation.

**Figure 4.**
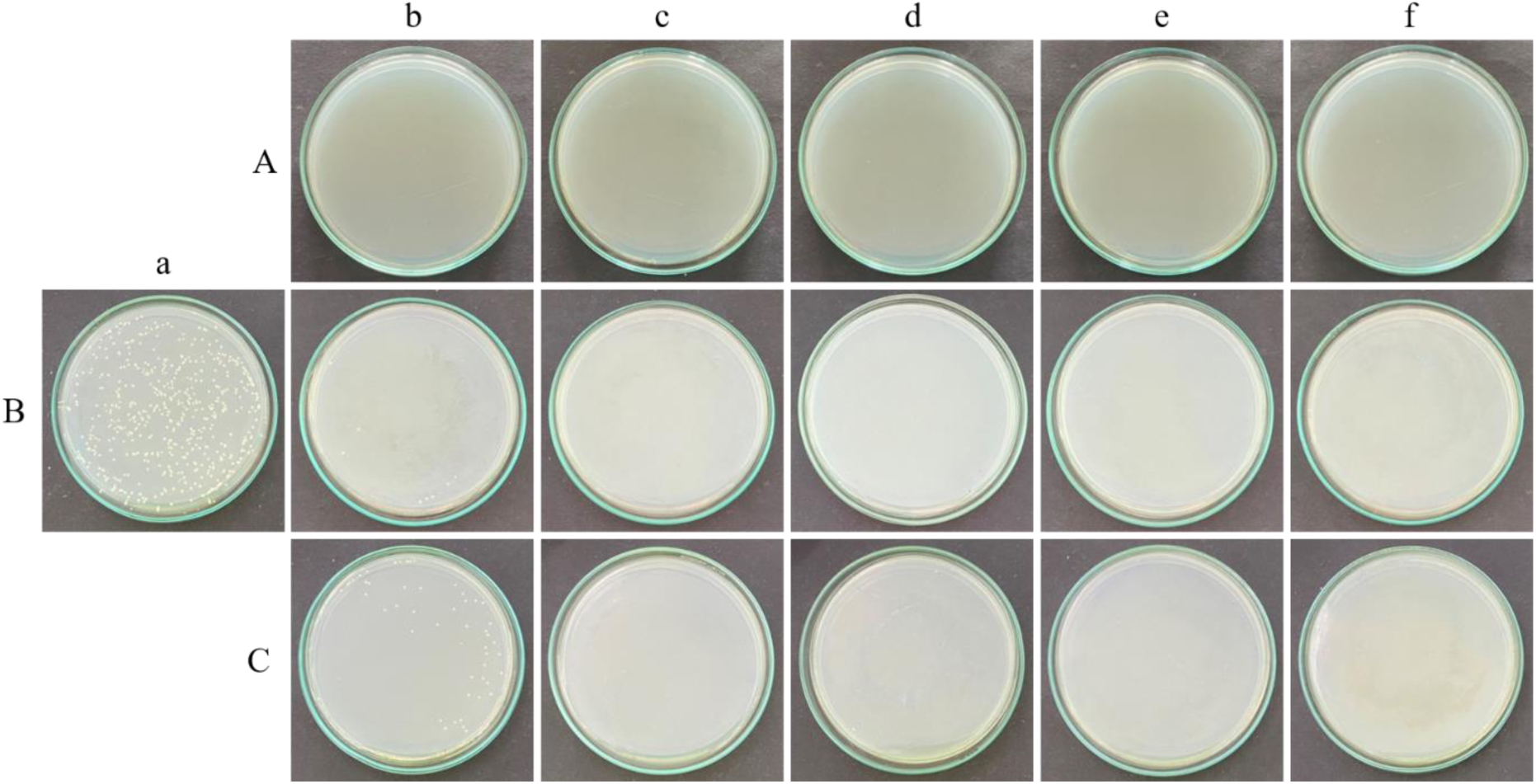
Effect of UV irradiation on *B. subtilis* assessed by CFU assay on nutrient agar plates. Bacterial suspensions were subjected to UV exposure at three distances from the light source: (A) 1 m, (B) 3 m, and (C) 5 m. For each distance, nutrient agar plates depict (a) untreated control cultures and those irradiated for (b) 15 min, (c) 30 min, (d) 45 min, (e) 60 min, and (f) 90 min. A noticeable decline in colony density with increasing exposure duration and closer proximity to the UV source highlights the time- and distance-dependent bactericidal activity of UV treatment against *B. subtilis*.

**Figure 5.**
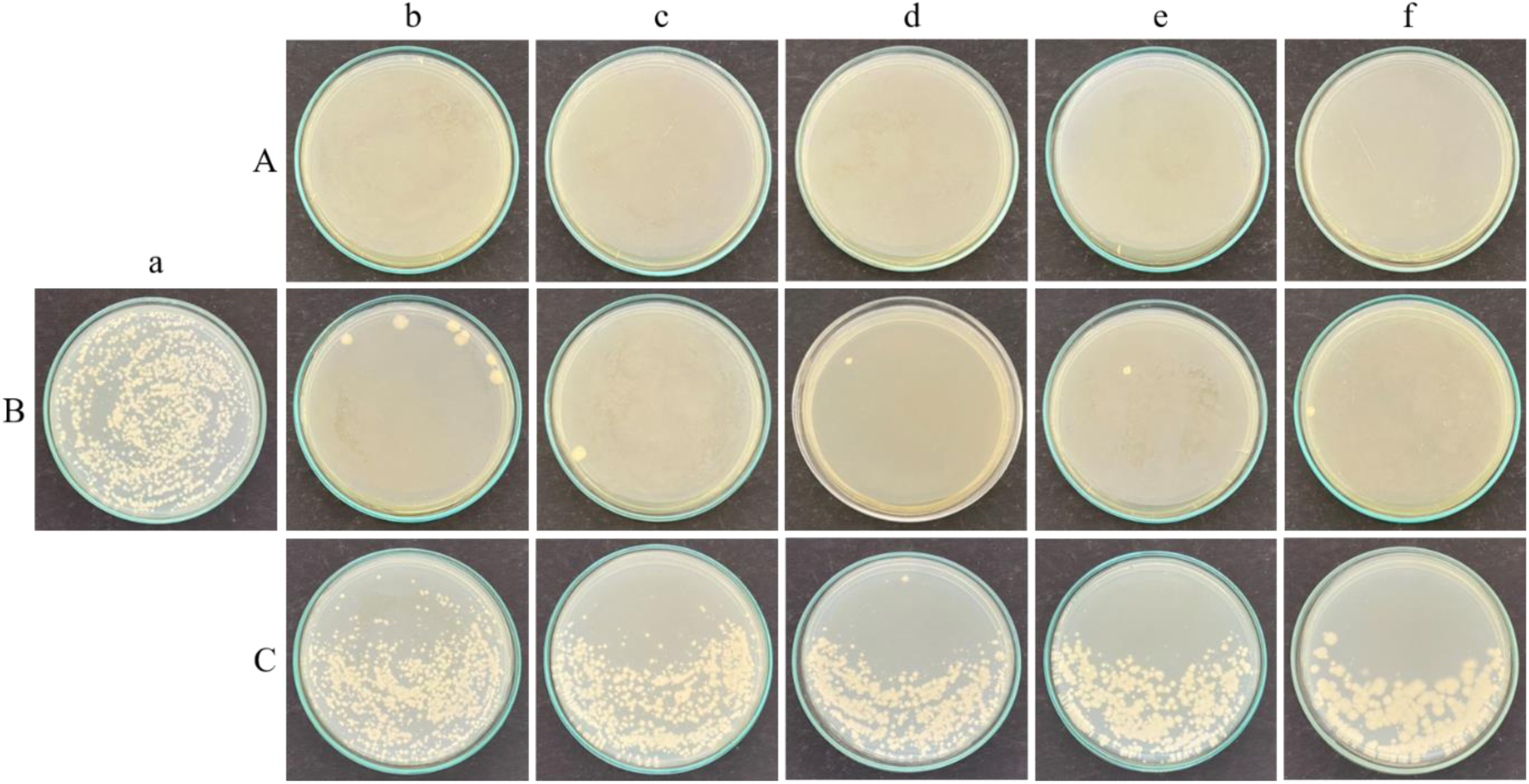
Effect of UV irradiation on *C. albicans* M-207 assessed by CFU assay on nutrient agar plates at (A) 1 m, (B) 3 m, (C) 5 m distances for (a) Control (b) 15 min (c) 30 min (d) 45 min (e) 60 min (f) 90 min in YEPD media. A decrease in colony count was observed with an increase in exposure time, and also the formation of crescent-shaped aggregates of colonies was observed at all time durations at a distance of 5 m.

**Figure 6.**
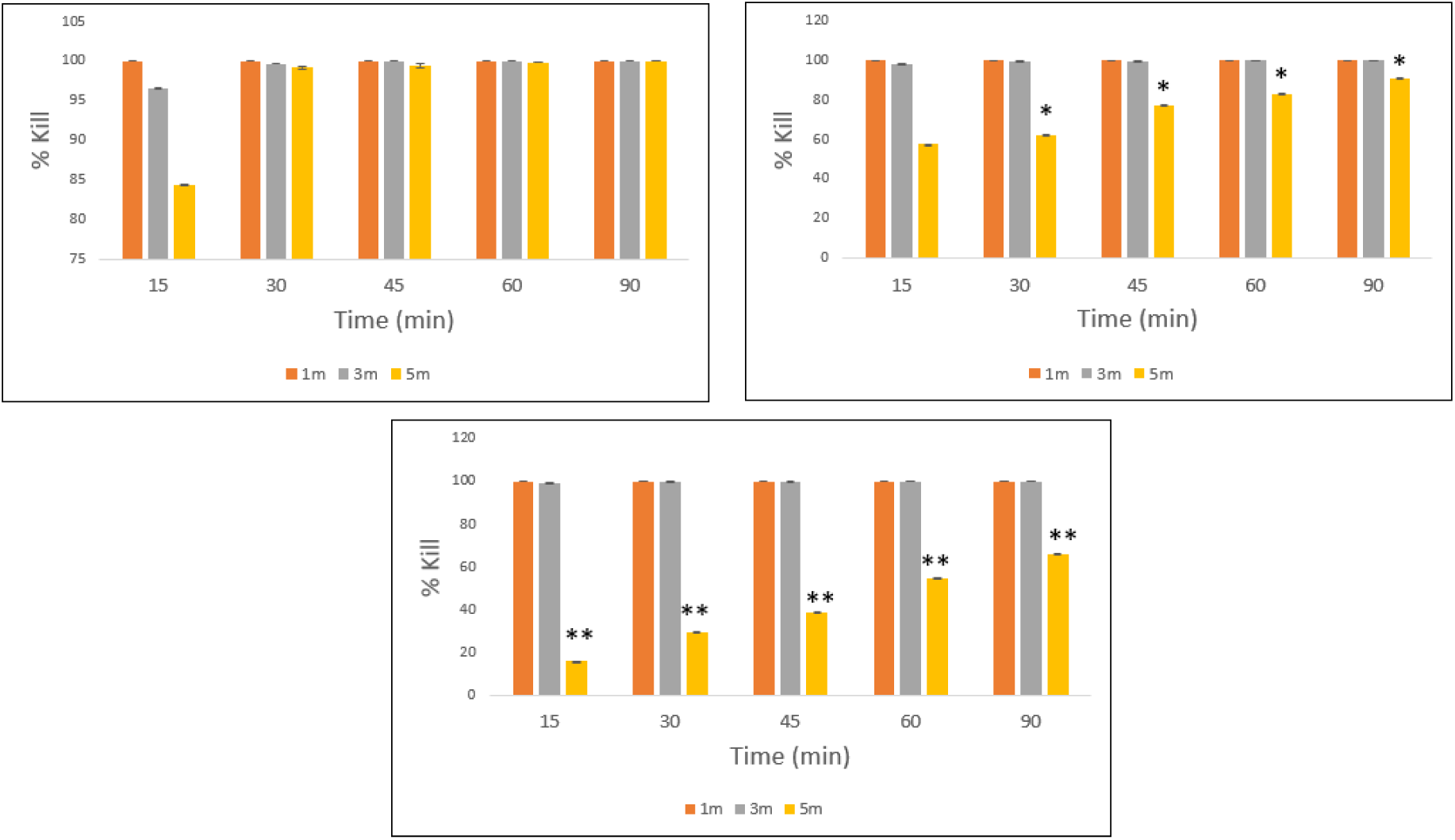
Percentage kill of (A) *B. subtilis*, (B) *E. coli* K12, (C) *C. albicans* M-207 on UV irradiation at different distances and time intervals. Asterisk represents the significant difference with respect to 1m distance with \**p* ≤ 0.05, \*\**p* ≤ 0.01.

**Table 1.**
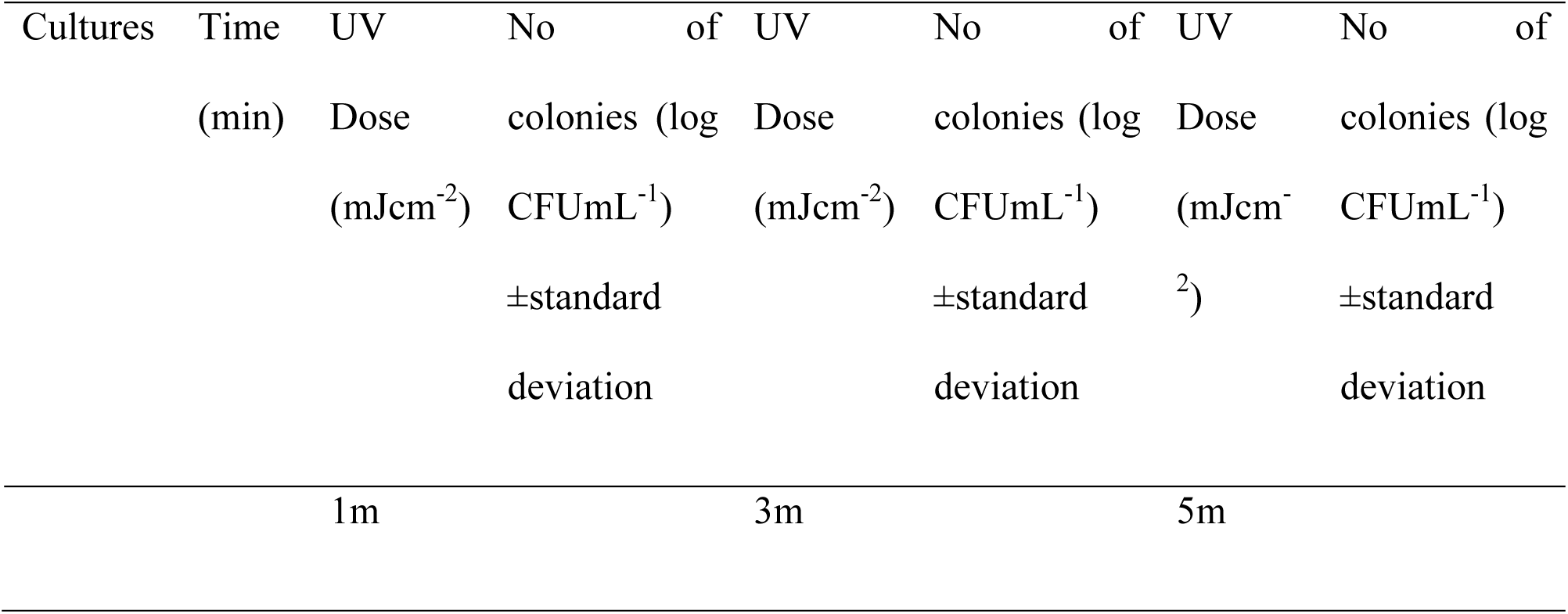
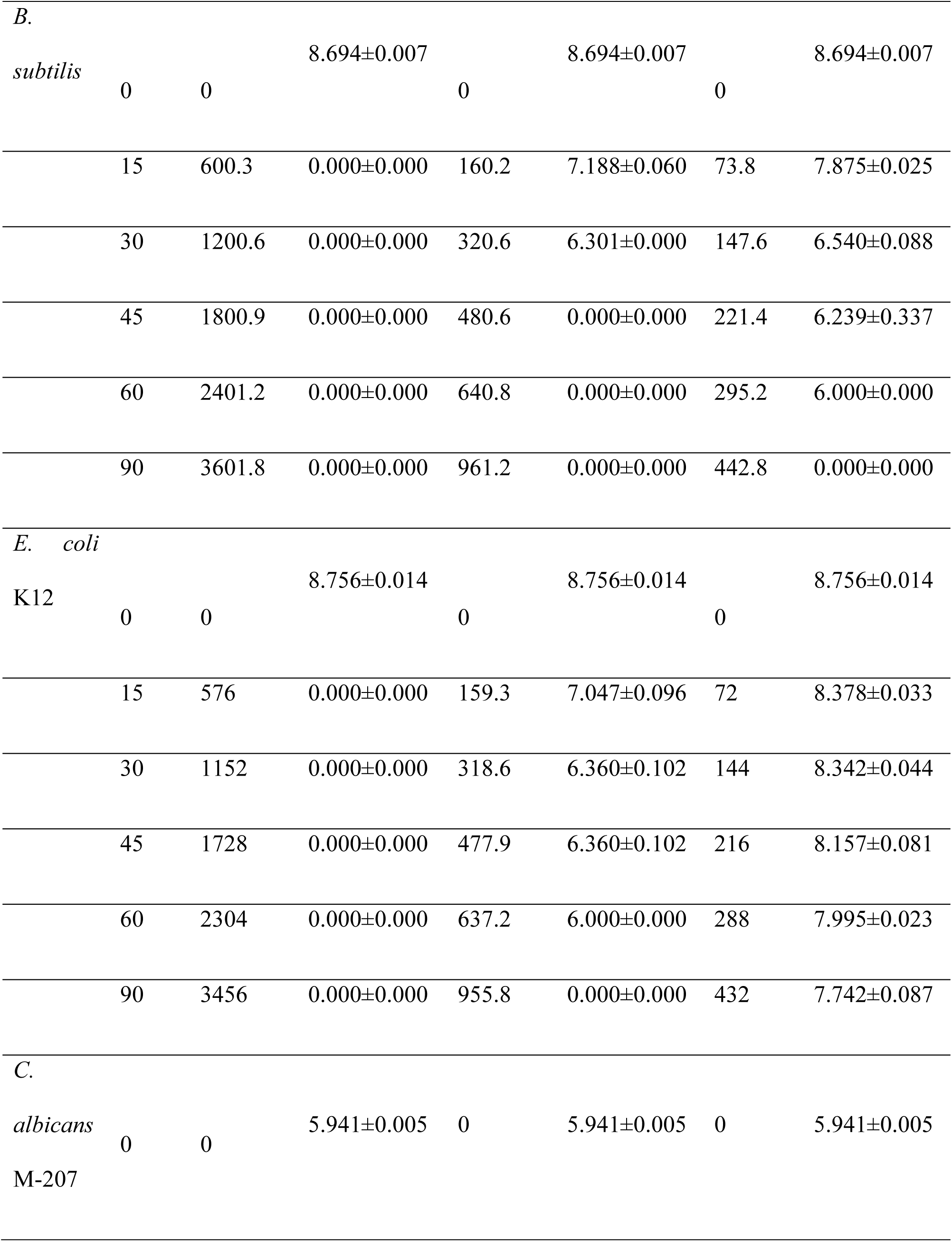
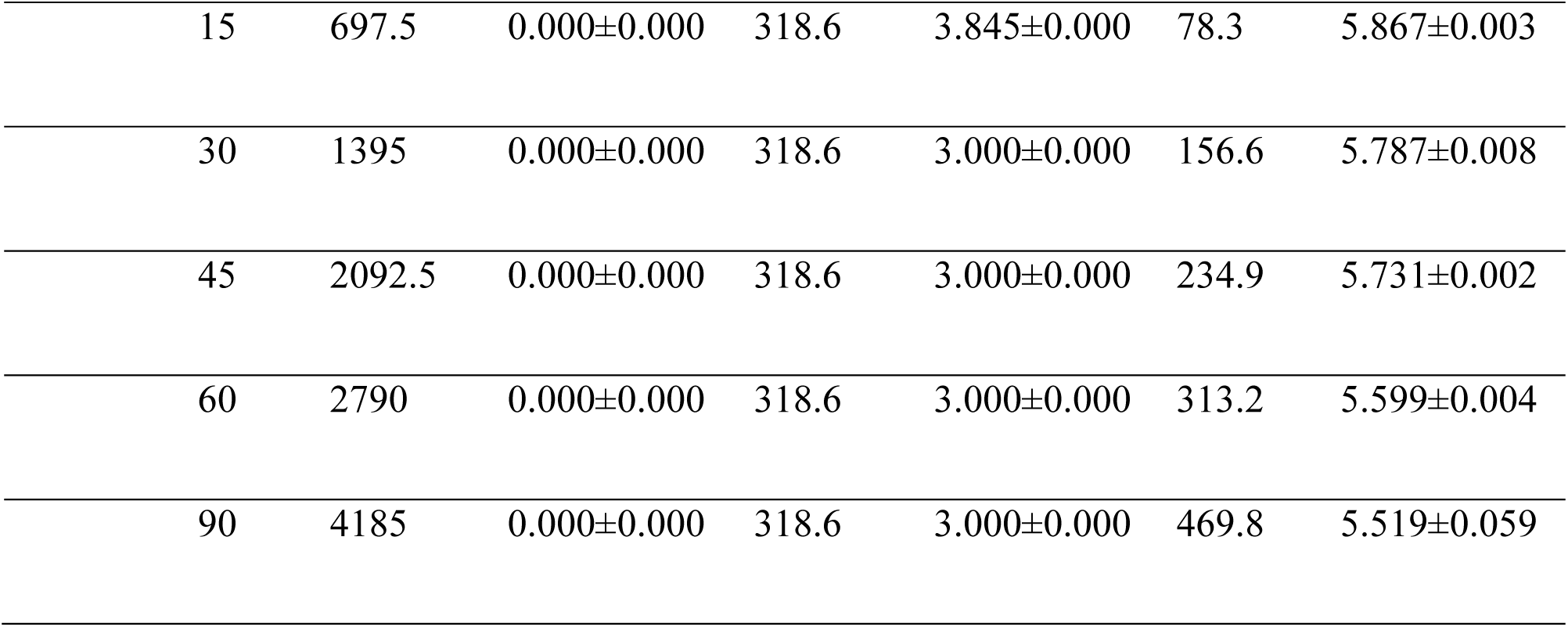
CFU count for *B. subtilis, E. coli* K12, and *C. albicans* M-207 at different time intervals and distances and the corresponding UV dose.

**Table 2.** Effect of UV exposure Distance, Time & Dose on Microbial cell viability & Inactivation (% kill).

| Organism | Exposure Distance (m) | Exposure Time (min) | UV Irradiance (mW/cm<sup>2</sup>) | UV Dose (mJ/cm<sup>2</sup>) | CFU- (% cell viability) | CFU- Inactivation (% kill) |
| --- | --- | --- | --- | --- | --- | --- |
| *B. subtilis* | 1 | 15 | 0.667 | 600.3 | 0 | 100 |
|  | 3 | 30 | 0.178 | 320.6 | 0.4 | 99.60 |
|  | 5 | 30 | 0.082 | 147.6 | 0.8 | 99.20 |
|  | 5 | 60 | 0.0820 | 295.2 | 0.20 | 99.80 |
|  | 5 | 90 | 0.0820 | 442.8 | 0 | 100 |
| *E. coli* K12 | 1 | 15 | 0.64.0 | 576.0 | 0 | 100 |
|  | 3 | 30 | 0.177 | 318.6 | 0.35 | 99.65 |
|  | 3 | 45 | 0.177 | 477.9 | 0.35 | 99.65 |
|  | 3 | 90 | 0.177 | 955.8 | 0 | 100 |
|  | 5 | 90 | 0.08.0 | 432.0 | 9.12 | 90.88 |
| *C. albicans*<br>*M-207* | 1 | 15 | 0.775 | 697.5 | 0 | 100 |
|  | 3 | 30 | 0.177 | 318.6 | 0.11 | 99.89 |
|  | 3 | 45 | 0.177 | 477.9 | 0.11 | 99.89 |
|  | 3 | 60 | 0.177 | 637.2 | 0.11 | 99.89 |
|  | 5 | 90 | 0.087 | 469.8 | 34.09 | 65.91 |

#### Viability of the cultures exposed to UVGI

The viability of the cultures *B. subtilis, E. coli* K12, and *C. albicans* M-207 on exposure to UV-C was determined using the MTT assay. In *B. subtilis,* the least cell viability of 78.81% was observed at 3 m for 30 min. In *E. coli* K12 at 3 m distances, low cell viability of 46.97% was observed when exposed to UV for 90 min. *C. albicans* M-207 exhibited 42.71% cell viability at 3 m for 60 min. The cell viability percentage and reduction in viable cells percentage of *B. subtilis*, *E. coli* K12, and *C. albicans* M-207 at different exposure distances and times is shown in Figure 7, Table 3 & 4. MTT analysis demonstrated a significant decrease in metabolic activity following UV exposure across all tested conditions. The reduction in cell viability was statistically significant compared to the control group (*p ≤ 0.05–0.0001*), depending on the exposure distance and duration.

**Figure 7.**
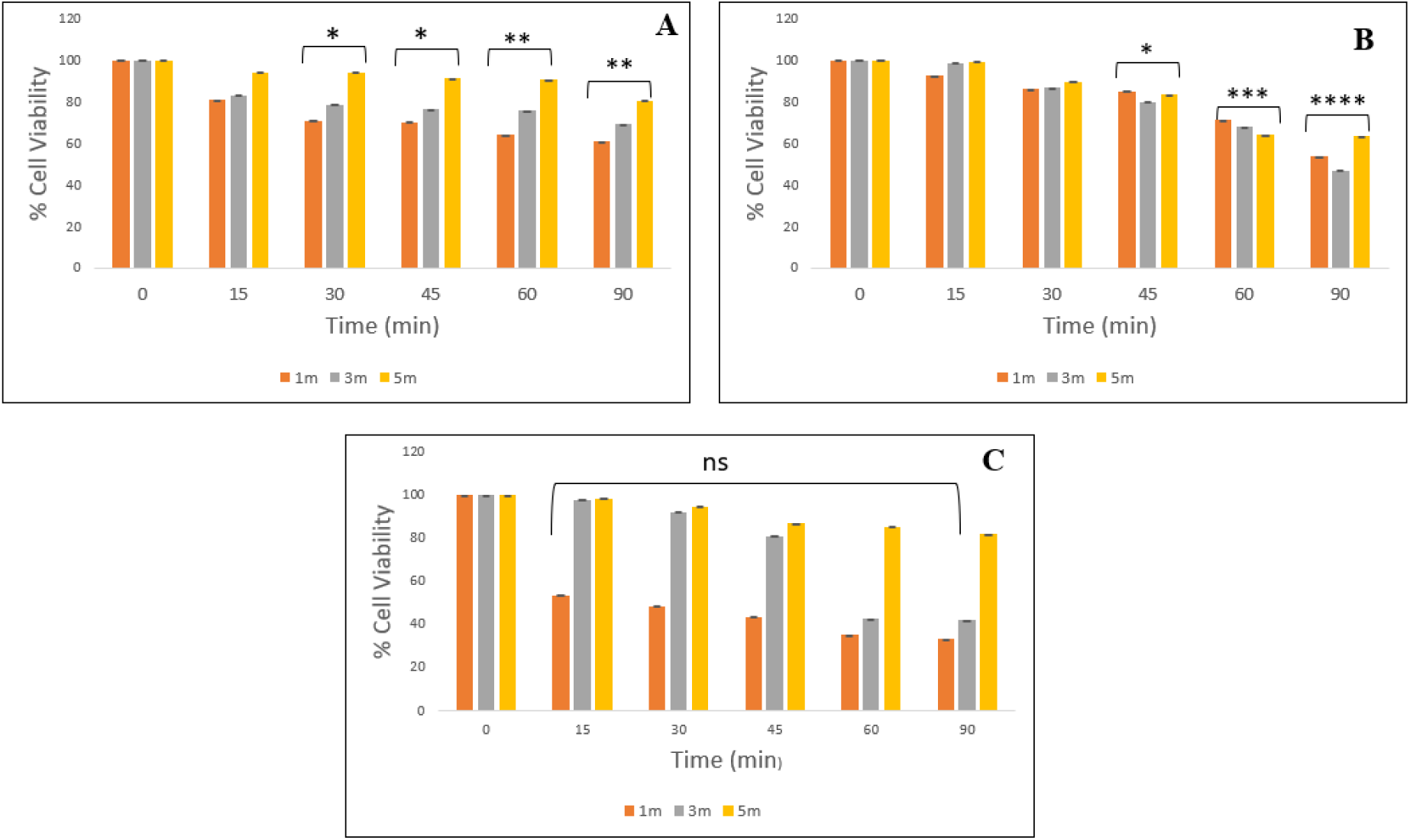
Cell viability analysis using MTT assay to validate the effect of UV exposure on (A) *B. subtilis,* (B) *E. coli* K12, (C) *C. albicans* M-207 at different distances and time intervals. Asterisk represents the significant difference with respect to the control with \**p* ≤ 0.05, \*\**p* ≤ 0.01, \*\*\**p* ≤ 0.001, \*\*\*\**p* ≤ 0.0001.

**Table 3.**
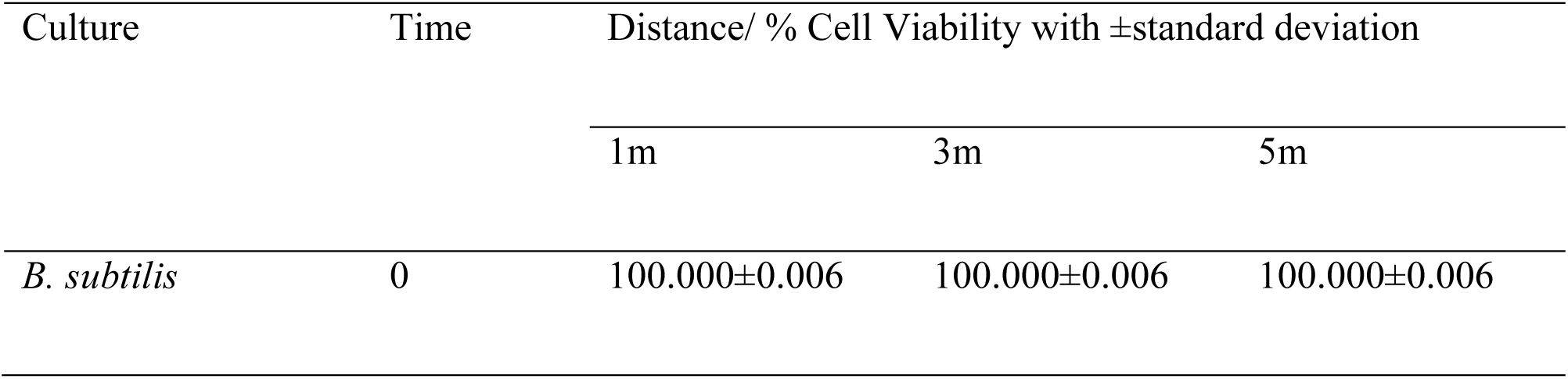
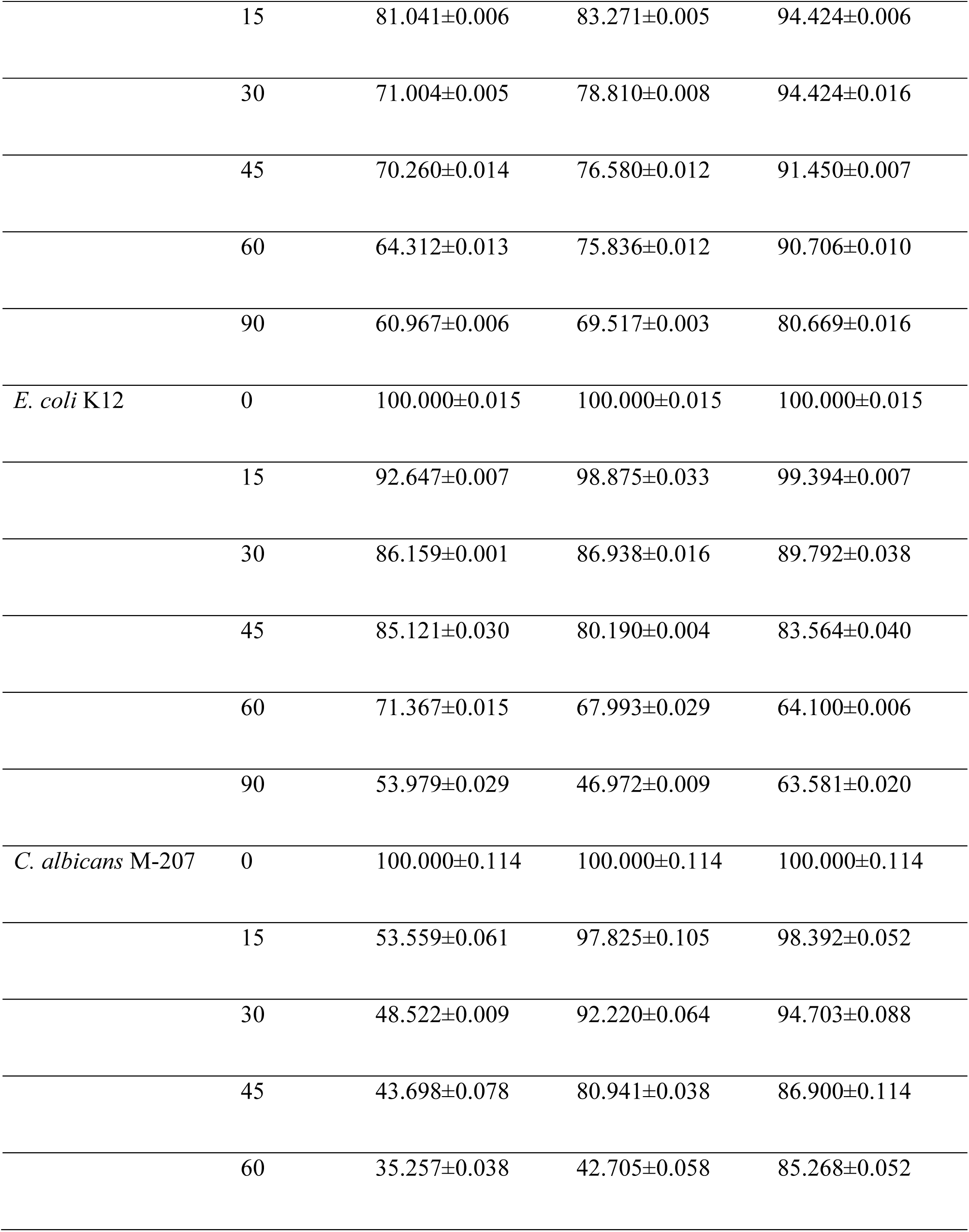
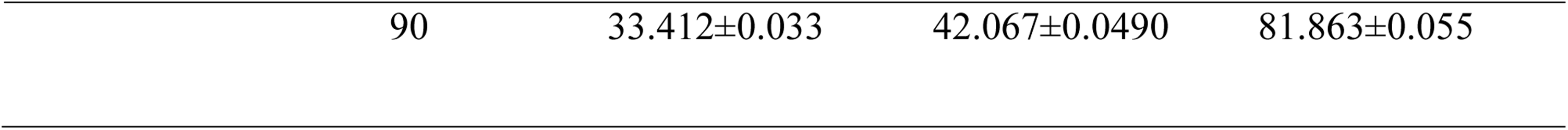
Cell viability of *B. subtilis*, *E. coli* K12 and *C. albicans* M-207 at different exposure distances and time.

| Culture | Time | Distance/ % Cell Viability with ±standard deviation | | |
| --- | --- | --- | --- | --- |
|  |  | 1m | 3m | 5m |
| *B. subtilis* | 0 | 100.000±0.006 | 100.000±0.006 | 100.000±0.006 |
|  | 15 | 81.041±0.006 | 83.271±0.005 | 94.424±0.006 |
|  | 30 | 71.004±0.005 | 78.810±0.008 | 94.424±0.016 |
|  | 45 | 70.260±0.014 | 76.580±0.012 | 91.450±0.007 |
|  | 60 | 64.312±0.013 | 75.836±0.012 | 90.706±0.010 |
|  | 90 | 60.967±0.006 | 69.517±0.003 | 80.669±0.016 |
| *E. coli* K12 | 0 | 100.000±0.015 | 100.000±0.015 | 100.000±0.015 |
|  | 15 | 92.647±0.007 | 98.875±0.033 | 99.394±0.007 |
|  | 30 | 86.159±0.001 | 86.938±0.016 | 89.792±0.038 |
|  | 45 | 85.121±0.030 | 80.190±0.004 | 83.564±0.040 |
|  | 60 | 71.367±0.015 | 67.993±0.029 | 64.100±0.006 |
|  | 90 | 53.979±0.029 | 46.972±0.009 | 63.581±0.020 |
| *C. albicans* M-207 | 0 | 100.000±0.114 | 100.000±0.114 | 100.000±0.114 |
|  | 15 | 53.559±0.061 | 97.825±0.105 | 98.392±0.052 |
|  | 30 | 48.522±0.009 | 92.220±0.064 | 94.703±0.088 |
|  | 45 | 43.698±0.078 | 80.941±0.038 | 86.900±0.114 |
|  | 60 | 35.257±0.038 | 42.705±0.058 | 85.268±0.052 |
|  | 90 | 33.412±0.033 | 42.067±0.0490 | 81.863±0.055 |

**Table 4.** Percentage reduction in viable cells of *B. subtilis*, *E. coli* K12, and *C. albicans* M-207 at different exposure distances and times.

| Culture | Time | Distance/ % kill with ± standard deviation |  |  |
| --- | --- | --- | --- | --- |
|  |  | 1m | 3m | 5m |
| *B. subtilis* | 0 | 0.000±0.006 | 0.000±0.006 | 0.000±0.006 |
|  | 15 | 18.959±0.006 | 16.729±0.005 | 5.576±0.006 |
|  | 30 | 28.996±0.005 | 21.190±0.008 | 5.576±0.016 |
|  | 45 | 29.740±0.014 | 23.420±0.012 | 8.550±0.007 |
|  | 60 | 35.688±0.013 | 24.164±0.012 | 9.294±0.010 |
|  | 90 | 39.033±0.006 | 30.483±0.003 | 19.331±0.016 |
| *E. coli* K12 | 0 | 0.000±0.015 | 0.000±0.015 | 0.000±0.015 |
|  | 15 | 7.353±0.007 | 1.125±0.033 | 0.606±0.007 |
|  | 30 | 13.841±0.001 | 13.062±0.016 | 10.208±0.038 |
|  | 45 | 14.879±0.030 | 19.810±0.004 | 16.436±0.040 |
|  | 60 | 28.633±0.015 | 32.007±0.029 | 35.900±0.006 |
|  | 90 | 46.021±0.029 | 53.028±0.009 | 36.419±0.020 |
| *C. albicans* M-207 | 0 | 0.000±0.114 | 0.000±0.114 | 0.000±0.114 |
|  | 15 | 46.441±0.061 | 2.175±0.105 | 1.608±0.052 |
|  | 30 | 51.487±0.009 | 7.780±0.064 | 5.297±0.088 |
|  | 45 | 56.302±0.078 | 19.059±0.038 | 13.100±0.114 |
|  | 60 | 64.743±0.038 | 57.295±0.058 | 14.732±0.052 |
|  | 90 | 66.588±0.033 | 57.933±0.0490 | 18.137±0.055 |

For *C. albicans* M-207, we observed 42.71% metabolic activity by MTT at 3 m for 60 min, while CFU results showed 0% viability at an exposure distance of 1 m for 15 min with an average UV dose of 697.5 mJ/cm². MTT and CFU assays evaluate different aspects of microbial viability and, therefore, may yield differing results following UVGI exposure. The MTT assay reflects short term metabolic activity, which can persist in cells that are damaged but not immediately inactivated. In contrast, CFU enumeration measures the ability of cells to proliferate and form colonies and is therefore considered a more stringent indicator of long-term viability.

In the present study, residual metabolic activity was detected in certain conditions despite a complete loss of culturability. This may be attributed to the presence of metabolically active but non-culturable cells, as well as persister subpopulations that retain enzymatic activity following stress exposure. Additionally, differences in assay sensitivity contribute to this variation, as MTT can detect low levels of cellular metabolism that do not translate into reproductive capacity. Therefore, CFU analysis represents a stricter endpoint for assessing antimicrobial efficacy, while MTT provides complementary information on short-term metabolic activity within the treated microbial population.

#### Scanning Electron Microscopy (SEM)

Structural characterization using SEM was performed to observe the surface morphology of cultures grown on silicone elastomer discs after UV-C exposure. When the control 5000X images of *B. subtilis, E. coli* K12, and *C. albicans* M-207 were compared, it was observed that *C. albicans* M-207 formed a very dense biofilm as compared to the other two cultures. Increased aggregation of cells was observed in *B. subtilis* when exposed to UV-C compared to the control. Spore formation in *B. subtilis* was observed at greater UVGI distances. In the case of *E. coli* K12, UVGI was the most effective at a 1 m distance when compared to 3 and 5 m. The dense biofilms of *C. albicans* M-207 were substantially reduced when exposed to the UVGI system for 30 and 60 min at distances of 1 and 3 m, respectively. However, UVGI was not effective in controlling biofilm formation at a distance of 5 m in *C. albicans* M-207 (Figure 8). The 1900x magnification images of the above-mentioned UVGI exposure studies are provided in the supplementary file (Supplementary Figure 1).

**Figure 8.**
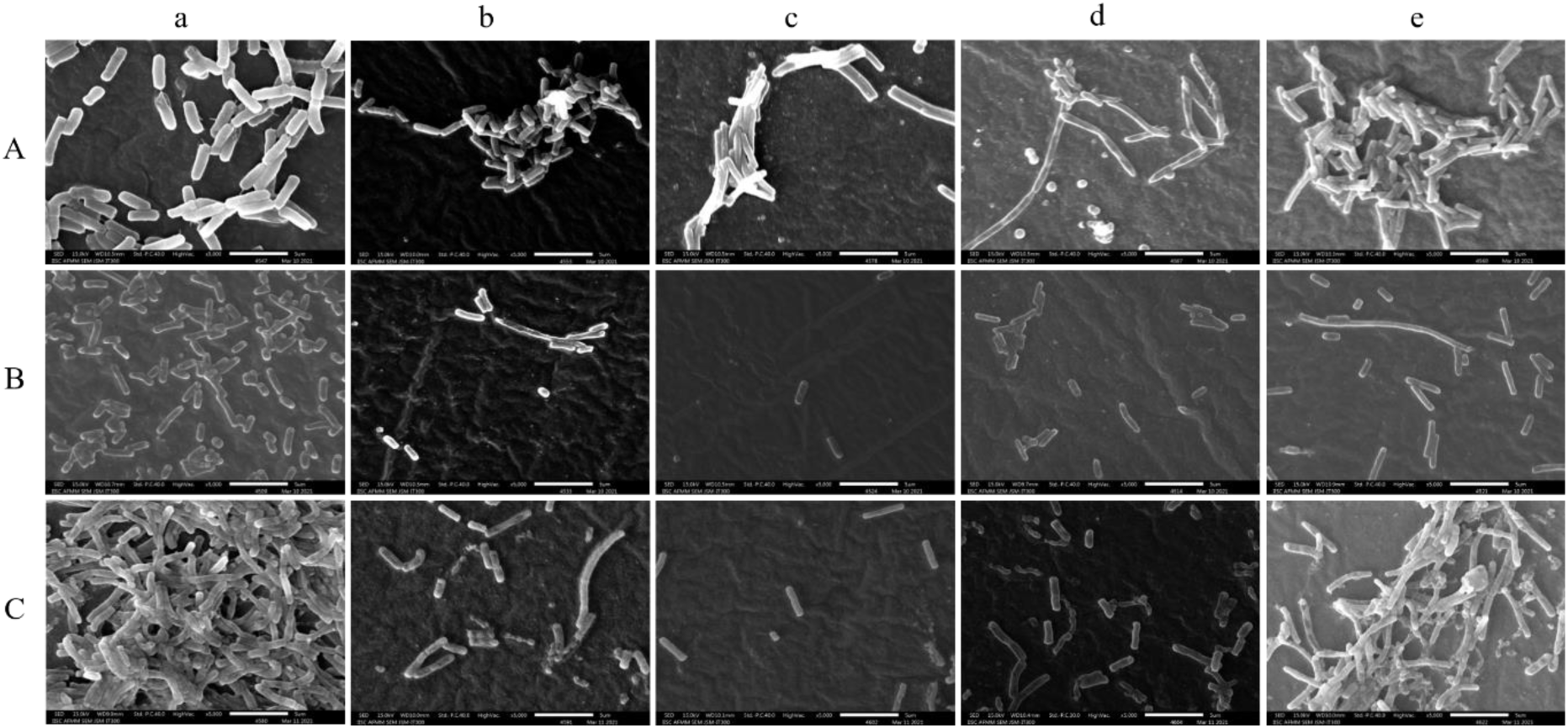
Scanning Electron Microscopy (SEM) images of representative biofilm-forming microorganisms: (A) *Bacillus subtilis*, (B) *Escherichia coli* K12, and (C) *Candida albicans* M-207 under different UV-C exposure conditions. For each organism, panels show: (a) UV C untreated control – cells are embedded in ECM and exhibit intact biofilm morphology; (b) Cells exposed to UV C at 1 m for 30 min – destruction of ECM, clumping of cells, loss of membrane integrity, and a pronounced amount of cell death; (c) Cells exposed to UV C at 1 m for 60 min – revealing cell membrane disruption and a reduction in the number of cells, indicating cell death; (d) Cells exposed to UV C at 3 m for 60 min – biofilm structure is disrupted, ECM is not visible, and cell contents are released; and (e) Cells exposed to UV C at 5 m for 60 min – reduction in ECM, but only a few cells are visible in the microscopic field, indicating cell death. Therefore, less distance and more exposure time lead to greater microbial killing. All SEM images were acquired at 5000× magnification.

SEM images captured at 1900× and 5000× magnifications revealed significant morphological alterations in *B. subtilis*, *E. coli* K12, and *C. albicans* M-207 following UV exposure. Control cells showed intact and typical cellular morphology, whereas treated cells exhibited surface irregularities, membrane damage, reduction in the ECM & number of cells, indicating cell death, deformation, and structural collapse. Higher magnification (5000×) highlighted detailed surface disruptions, while lower magnification (1900×) showed reduced cell density and aggregation changes.

## DISCUSSION

Infections are classified as hospital-acquired if they occur within 48-72 h of the visit to the hospital or if they appear within 10 days after hospital discharge (WHO, 2002). Hospital-acquired infections are caused by many microorganisms, predominantly gram-positive and gram-negative bacteria, fungi, and viruses (25). *E. coli* is one of the bacteria that must be tested for household/hospital disinfection. A compact, in-house, easy-to-use, and low-cost UVGI sterilization system was developed in the current study. This system was evaluated for targeted sterilization of hospital wards and operating theatres, which was validated using model bacterial and fungal systems. However, this system can also be used in various other places, such as schools, malls, restaurants, theatres, banks, shopping centres, airports, airplanes, public toilets, and public transport. With little modification to the existing manual portable UVGI system, wall-mounted, table top, and automated portable models can also be developed. Bacteria and fungi form community structures known as biofilms that are difficult to control. These biofilms are composed of microbial cells, proteins, carbohydrates, and enzymes in an extracellular matrix (ECM). There are channels within the ECM for the transport of nutrients, antimicrobials, waste, etc. There are also void spaces in which nutrients, waste, and antimicrobials are stored. Biofilms formed by these microorganisms protect them by absorbing UV, scavenging ROS, increasing the path length, and scattering UV (7). Studies have used UV-C radiation on biofilms and have successfully controlled these biofilms (7, 26). However, additional treatments may be required to thoroughly remove the biofilms (8). If not thoroughly removed, new growth of organisms may occur on the remains of the previous biofilm (27).

Mitigation of microbial contamination and its associated infections by direct exposure to UV irradiation is being practiced as a chemical-free approach or in tandem with other sanitation measures to maintain a sterile environment (28, 29). Therefore, UV radiation is a proven technique for chemically free sterilization. However, their effects on biofilm-forming microorganisms are limited. Hypothetically, UV radiation inactivates cells within the biofilm, thus slowing down the process of biofilm formation (7). The continuous/pulsed mode of UV irradiation has proven to be effective in sterilization (30, 31) and reduced biofilm formation (8).

The UVGI system was effective in controlling biofilm-forming bacteria and yeast. The ECM in the biofilm is damaged by UV-C, causing the disruption and leakage of components. Due to damage to the biofilm ECM, UV-C comes in direct contact with the cell wall, leading to cell damage and death. UV-C photons are also absorbed by intracellular components such as proteins, DNA, and RNA, causing their denaturation/damage. Damage to nucleic acids can occur in the form of double bonds/dimers between adjacent nucleotides. This photochemical damage is known as pyrimidine dimerization, the most common of which is thymine dimerization (Figure 9).

**Figure 9.**
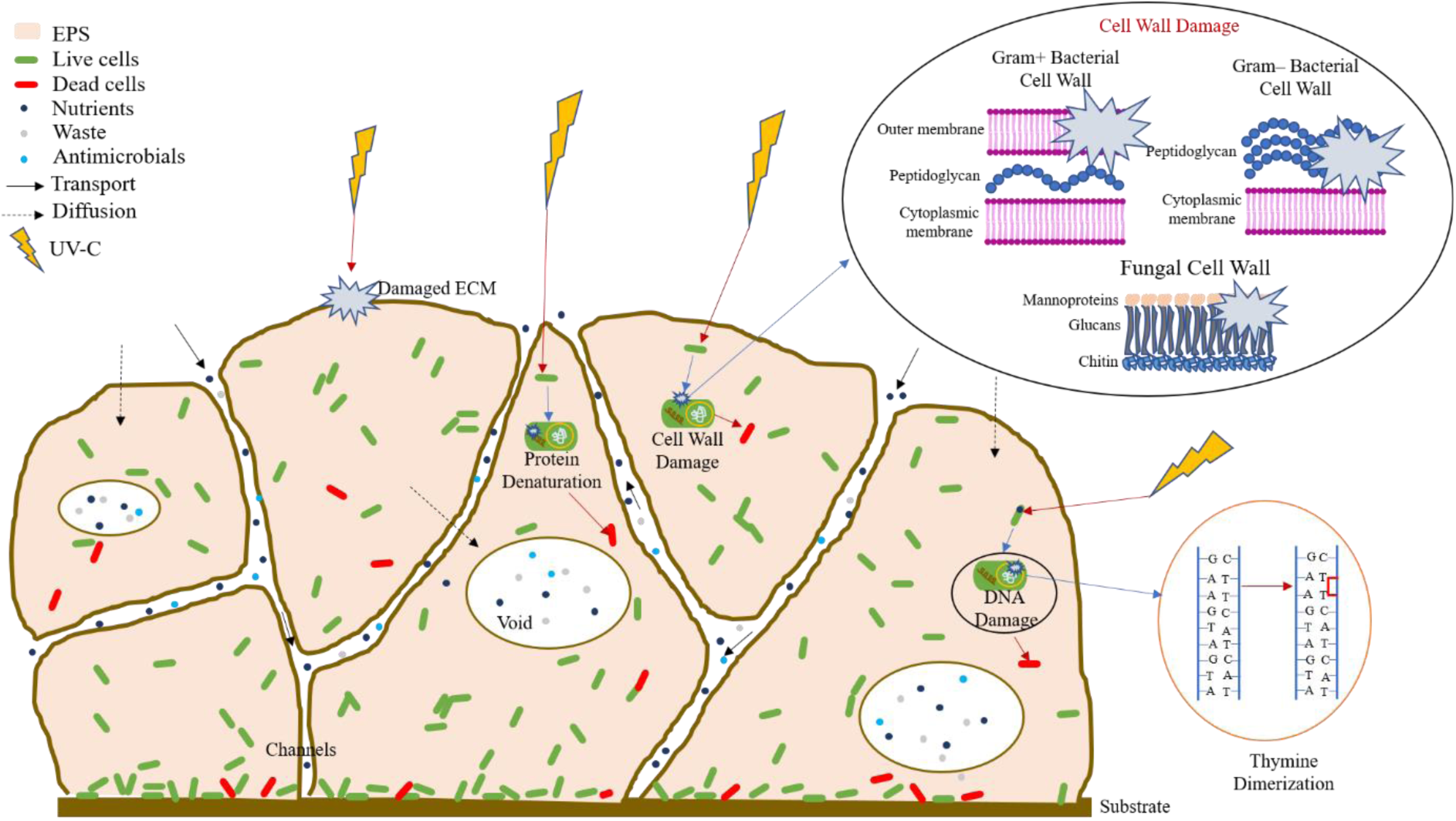
Schematic representation of the microbicidal mechanism of action of UV-C on bacteria and fungi.

These three microorganisms - *Bacillus subtilis* (Gram-positive, spore-forming, motile bacterium), *Escherichia coli* K12 (Gram-negative, non-spore-forming, non-motile bacterium), and *Candida albicans* M-207 (multi-drug resistant, clinical yeast isolate)- were selected because they are all robust biofilm formers and represent different microbial categories commonly encountered in real environments. *B. subtilis* is ubiquitous, can grow in aerobic as well as in anaerobic conditions, found in soil, water, air, dust, plants, foods, and hospital settings, and can cause secondary nosocomial infections, including wound, respiratory, and systemic infections. *E. coli* K12, an enteric bacterium belonging to the ESKAPEE group (E – *Enterococcus faecium*, S – *Staphylococcus aureus*, K – *Klebsiella pneumoniae*, A – *Acinetobacter baumannii*, P – *Pseudomonas aeruginosa,* E – *Enterobacter* species, E- *Escherichia coli*), represents Gram-negative pathogens commonly involved in environmental contamination and clinical infections. *C. albicans* represents clinically relevant fungal biofilms that are resistant to multiple drugs. Together, these models provide a relevant spectrum of biofilm-forming organisms for evaluating UVGI disinfection, mimicking real-world microbial contamination in hospitals, public spaces, and environmental niches. Three different models were used to study the germicidal action of the UVGI system: a Petri plate model to assess viable count (CFU), 96 well microtiter plate model to assess cell viability (MTT), and a silicone elastomer disc model for morphological analysis of the test organisms (using SEM).

In this study, sensitivity to UV-C was observed for *B. subtilis*, *E. coli* K12, and *C. albicans* M-207 at different time intervals and distances with cell killing/inhibition observed when CFU were counted. A linear reduction in the number of cells with respect to dose and time for all cultures was observed for bacteria, similar to a published study (31). However, in the case of *C. albicans* M-207 reduction was observed, even at a lower intensity of UV irradiation, as the exposure time increased. Although there was a definite reduction in colony count, the increase in colony size was striking. Whenever there are fewer colonies, the colony size is large because there is minimum competition for space and nutrients (crowded plates have small colonies). We also observed an interesting phenomenon in *C. albicans* M-207, where at low UV-C, the colonies tended to aggregate in a crescent shape towards one half of the petri dish facing the radiation appearing partially inhibited. However, this phenomenon has not yet been observed in other cultures. The Petri plate side directly exposed to UV-C formed this crescent-shaped aggregate, whereas the other side had no colonies. This is because of the petri plate barrier, the UV-C cannot penetrate that side, and the colonies are shielded from the radiation, whereas the colonies on the other side are killed. This also demonstrates that UV-C radiation cannot penetrate glass and polymeric materials, such as polystyrene. The most refractory material for UV-C disinfection is plastic, followed by glass, and stainless steel (32). A reduction in cell viability was also observed as compared to the control, indicating the sensitivity of the cultures to UVGI, as reported previously (33) in another study.

SEM was performed as a qualitative method to evaluate cell death and understand the morphological changes in the microbes when treated with different doses of UV-C radiation at different distances (34, 35). SEM analysis revealed clear evidence of UV-induced structural damage, including disruption of cell surfaces, reduction in extracellular matrix (ECM), and decreased cell density across all tested microorganisms. These observations indicate progressive deterioration of biofilm architecture with increasing UV exposure. In the case of *C. albicans* M-207, residual biofilm-like structures were observed under certain exposure conditions, particularly at increased distances from the UV source. This likely reflects incomplete inactivation of the microbial population at lower UV doses. Such observations may be associated with the survival of stress-tolerant or partially damaged cells (36). Overall, the SEM findings demonstrate that reduced distance and increased exposure time enhance the extent of structural disruption, highlighting the importance of optimizing UV-C dose and exposure conditions for effective microbial inactivation.

The findings of this study align with recent reports demonstrating that UV-C irradiation is a promising strategy for inactivating biofilm-associated microorganisms, although efficacy strongly depends on microbial type, biofilm maturity, and exposure conditions. A study by (37) showed that at 254 nm UV-C irradiation was effective in reducing biofilms of healthcare-associated pathogens on stainless steel surfaces, supporting our observations of biofilm susceptibility under controlled UVGI treatment. Similarly, (38, 39) reported that *E. coli* biofilms displayed variable UV sensitivity depending on growth temperature and biofilm maturation. In line with our data on *C. albicans* (40) demonstrated that portable UV-C devices significantly disrupted *C. albicans* biofilms, highlighting the clinical relevance of targeting fungal biofilms with UV technologies. Furthermore, a recent critical review emphasized that biofilm extracellular matrix composition and heterogeneity often limit complete eradication, as metabolic assays (e.g., MTT) may overestimate viability compared to CFU counts. Collectively, these findings underscore that while UV-C treatment is broadly effective, discrepancies between viability assays reflect complex biofilm physiology and reinforce the value of combining metabolic and culture-based methods, as applied in our study.

Ultraviolet-C (UVC) light, with wavelengths ranging from 200–280 nm, has been widely studied for its germicidal properties and is effective against bacteria, fungi, and viruses by causing DNA/RNA damage. Conventional UVC systems, typically operating at 254 nm, are effective for air, surface, and water disinfection; however, direct exposure can be harmful to human skin and eyes, limiting their use in occupied spaces. UVC disinfection systems can be classified as near-UVC (conventional 254 nm) and Far-UVC (207–222 nm), with differences in microbial inactivation efficiency, penetration, and safety profiles. Emerging Far-UVC technology (207–222 nm) has gained attention as a potentially safer alternative. Studies suggest that Far-UVC can inactivate microbes efficiently while penetrating only the outer dead layer of human skin and the tear layer of the eyes, potentially reducing health risks associated with conventional UVC (41, 42, 43). While Far-UVC offers a novel approach for continuous disinfection in contaminated public spaces, issues such as lamp cost, dose optimization, and long-term safety must be addressed.

For effective biofilm disruption, especially in applications requiring deep penetration, conventional UVC (254 nm) is currently the more suitable choice. However, Far-UVC (222 nm) holds promise for applications in occupied spaces due to its safety profile, though its efficacy in biofilm disruption is still under investigation (44, 45). Therefore, our study focused on conventional UVC (253.7 nm) due to its well-characterized efficacy and safety profile under laboratory-controlled conditions, but future work could explore Far-UVC applications as the technology matures.

Current literature reporting UV-C disinfection of biofilms has been systematically compared with our results, and the outcomes are presented in Table 5. Our results, in line with these reports, indicate that biofilms show higher resistance compared to planktonic cells due to the protective EPS matrix. Importantly, the UV dose and exposure time determined whether only surface-associated cells were inactivated or whether deeper biofilm layers were also disrupted. Lower doses mainly reduced surface viability, while higher doses promoted more uniform microbial killing and disruption of the biofilm matrix. These findings emphasize that effective biofilm disinfection requires delivering sufficient UV dose to not only kill embedded cells but also overcome EPS-mediated protection, which is critical for the design of portable UVGI systems for real-world applications.

**Table 5.** Comparative Studies on UVGI Efficacy Against Biofilm-Forming Microorganisms with Existing Literature.

| Organism(s) | Inoculum density & Material | UV Source Type & Mobility | UV Dose (mJ/cm<sup>2</sup>), distance and time of exposure | CFU (Long-term viability assessment) | MTT (Metabolic activity/ Short-term viability assessment) | References |
| --- | --- | --- | --- | --- | --- | --- |
| *B. subtilis*<br>*E. coli* K12<br>*C.albicans*<br>M-207 | 1×10<sup>6</sup> cells/mL;<br>biofilm formed on petriplate, 96 well microtiter plate & silicone elastomer discs | UV tubes (Philips - 36W) UV-C (253.7 nm)<br>wireless, portable, laptop/mobile-connectable | 576 (1 m), 318.6(3m), 172.8 (5m)<br>1–5 m, 15–90 min | 1 m/15 min → 100% kill (all 3 cultures) | *B. subtilis* -64% viability (1 m, 60–90 min);<br>*E. coli* 47% (1 m, 90 min);<br>*C. albicans* <50% (1 m all intervals, 3 m at 60–90 min) | Present study |
| *S. aureus*,<br>*E. coli*,<br>*P.aeruginos*<br>*a*,<br>*C.albicans* | Biofilm<br>formed on<br>stainless<br>steel discs | UV-C<br>(254 nm)<br>Low-pressure<br>mercury<br>lamp;<br>wired;<br>bench-top<br>unit | 467.8–946.7 | >3 log<sub>10</sub><br>reduction in<br>bacterial<br>biofilms; *C.*<br>*albicans* - 3.17<br>log reduction | Not reported | (37) |
| *E. coli*<br>biofilms<br>(different<br>maturation:<br>1d @ 37 °C,<br>3d @ 25 °C,<br>5d @ 15<br>°C) | Biofilms<br>matured at<br>different<br>temps,<br>~10<sup>6</sup><br>CFU/cm<sup>2</sup> | UV-C<br>(254 nm),<br>Low-pressure<br>mercury<br>lamp;<br>wired; lab<br>chamber | ~30–180s<br>(lamp output<br>dependent) | Young<br>biofilms → 3.5<br>log reduction;<br>Mature<br>biofilms → 1.8<br>log reduction | Not reported | (38) |
| *Clostridioides difficile* spores | 10<sup>5</sup>–10<sup>7</sup> CFU/mL on agar plates & suspensions | UV-C (254 nm), UV-C lamp, fixed, wired; bench-top device | ~2208 (for sporicidal effect) up to 30 min | Complete inactivation after 20 min at high dose; spores highly resistant | Not reported | (39) |
| *Candida albicans* biofilm | Inoculum ~2.39×10<sup>6</sup> CFU/mL<br>Biofilm formed on PMMA (polymethyl methacrylate)(denture base) | UV-C Hand-held UV-C lamp; wired, portable but not wireless | NA (lamp-based) | Reduced from 2.39×10<sup>6</sup> to 2.15×10<sup>5</sup> CFU/mL (~91% kill) | Not reported | (40) |
| *Pseudomonas aeruginosa* | Biofilms grown in CDC reactor<br>Biofilms on Acrylonitrile Butadiene Styrene (ABS-thermoplastic polymer), High-Density Polyethylene (HDPE), | UV-LED 280 nm, doses varied by surface<br>wired; not mobile | NA (surface-dependent) | Log reductions depended on material:<br>steel/PC → higher kill; silicone/HDPE → more resistant | Not reported | (41) |
|  | Polycarbonate (PC),<br>steel,<br>silicone |  |  |  |  |  |

A limitation of UV disinfection lies in its restricted penetration into complex, multilayered biofilms. The extracellular polymeric substance (EPS) matrix, together with cell density and surface irregularities, attenuates UV radiation and shields embedded cells, allowing some viable microbes to persist despite surface inactivation. This structural protection implies that UV alone may not always ensure complete eradication of mature biofilms, especially in clinical or environmental settings where biofilms can exceed several hundred microns in thickness. Consequently, integrated or sequential approaches may be required. For example, combining UV with chemical disinfectants, photocatalytic agents, or mechanical disruption can enhance matrix breakdown and improve microbial inactivation. Such combined methods may lower the overall energy demand of UV while ensuring more complete biofilm eradication. Our findings underscore the importance of considering these synergistic strategies when developing portable UV disinfection systems for real-world contaminated environments.

The present study demonstrates that UVC irradiation can achieve effective inactivation of biofilm forming bacteria & fungi on different laboratory model systems, including petri plates, 96-well microtiter plates, and silicone elastomer discs. Although *in vitro* models inherently face limitations such as light scattering, absorption by plastic surfaces, and the presence of shadowed regions that may reduce UV penetration, our optimized experimental setup ensured uniform exposure and achieved 100% microbial inactivation under controlled conditions.

Importantly, the use of silicone elastomer discs, a material commonly employed in medical devices such as catheters, prosthetics, and implantable surfaces, enhances the clinical relevance of our findings. Since these materials are prone to biofilm-associated contamination in healthcare settings, demonstrating effective UV disinfection on this substrate provides valuable translational insight into potential real-world applications. However, it is important to recognize that real-world environments present additional challenges such as surface irregularities, complex geometries, shadowed zones, and heterogeneous organic loads, which can all limit UV penetration and efficacy. UVC penetration is limited, combined strategies such as mechanical cleaning, chemical disinfectants, or photocatalysis may be required to ensure complete removal of resilient biofilms.

We have used petridish, 96-well microtiter plate and silicone elastomer experimental models to study the Shielding/ barrier effect, shadow zone during UVGI exposure in the laboratory & found that at 1m, 15m exposure corresponding to doses of 600.3, 576 & 697.5 mJ/cm² is effective in the 100% kill of *B. subtilis*, *E. coli* and *C. albicans*. The limited penetration of UV-C through glass and most plastics such as polystyrene has important implications for the design and effectiveness of UVGI systems. Since germicidal UV-C is largely absorbed by these materials, microorganisms shielded behind barriers or embedded in multilayer biofilms may not be effectively inactivated. Recent studies highlight this challenge and potential solutions: for example, thin polystyrene surfaces allow limited UVC transmission while maintaining structural integrity and chamber designs incorporating reflective surfaces (46) and UV-transparent materials can significantly improve dose uniformity and microbial inactivation (47, 48). Modern reviews also emphasize the importance of accounting for optical losses and shielding effects when designing UV-LED and UV-C lamp systems for practical applications (49). These findings underscore that careful selection of materials, exposure geometry, and reflective surfaces is essential to translate laboratory results into reliable, real-world UVGI applications in healthcare, water treatment, and environmental sanitation.

Infectious disease outbreaks have led to an understanding of the importance of cleanliness, disinfection, and sterilization. Microbes are important and fundamental forms of life from the early stages of Earth’s evolution and therefore can survive under any environmental conditions and acclimatize due to adaptive mutations. Therefore, this study was conducted to determine the effectiveness of UVGI as a microbicidal platform. The UVGI disinfectant system was developed indigenously and is easy to operate, transport, and economical. It has been shown to effectively sterilize bacterial and fungal cultures, with a direct correlation between the exposure distance and time. Therefore, this system can be used for effective disinfection or sterilization of medical devices and hospital areas, such as waiting rooms, laboratories, operation theatres, isolation rooms, emergency rooms, schools, malls, restaurants, homes, and the food industry. The developed UVGI system can be scaled up, modified, and programmed according to requirements to ensure minimum infection risk.

## CONCLUSION

A compact, in-house indigenously developed UVGI disinfection system was successfully designed and validated as an effective microbicidal platform against biofilm-forming bacterial and fungal pathogens. The system is easy to operate, transport, and cost-effective, making it suitable for diverse disinfection applications. The UVGI system achieved complete inactivation of representative biofilm-forming bacteria (*B. subtilis*, *E. coli K12*) and fungi (*C. albicans M207*) under optimized conditions. Experimental studies such as CFU, MTT assay and SEM were conducted. MTT and CFU assays provide complementary insights into long-term viability and short-term metabolic activity, respectively. CFU assay demonstrated 100% kill of all tested organisms at 1 m and 15 min, corresponding to doses of 600.3, 576 & 697.5 mJ/cm². Morphological analyses (SEM) confirmed significant biofilm disruption, ECM destruction, and cell death, validating the system’s efficacy against resilient microbial communities. A dose & time-dependent inactivation of biofilm-forming bacteria & fungi was observed on exposure to UVGI. The device also shows strong potential for real-world deployment in high-risk environments, including medical facilities (operating theatres, isolation rooms), public spaces (schools, malls, restaurants), homes, and the food industry. The modular design of this device also allows customization and automation, enabling integration into mobile or robotic disinfection platforms for enhanced infection control. Therefore, the findings demonstrate proof-of-concept efficacy of the developed UVGI system against biofilm-forming microorganisms under controlled laboratory conditions and suggest its potential for further validation in clinical environments.

## CRediT authorship contribution statement

**Bindu Sadanandan**: Conceptualization, Methodology, Validation, Formal analysis, Investigation, Data curation, Writing - Review and Editing, Visualization, Supervision. **Shyam Sunder**: Conceptualization, Software, Visualization, Supervision. **Vaniyamparambath Vijayalakshmi**: Formal analysis, Investigation, Data curation, Writing – Original draft. **Priya Ashrit**: Investigation, Data curation, Writing – Original draft. **Kavyasree Marabanahalli Yogendraiah**: Formal analysis, Investigation, Data curation, Writing – Original draft. **Kalidas Shetty**: Writing - Review and Editing.

## Acknowledgement

The authors wish to thank the Director R&D, General Manager (PD & IC), and Chief Technical Officer (EO & L) for providing the necessary facilities at BEL Limited. We also thank Charu S Tripati, Sr. DGM (EO & L) & Damodar Kadaba, Sr. MRS (CRL), for continuous motivation. We would also like to thank Durga Rao (Energy Systems, PDIC) for providing valuable reviews on the power supply of the system. We appreciate the effort put in by the Contract Engineers, Ashwitha and Babu in the assembly of the system.

## Data Availability Statement

The data that support the findings of this study are available from the corresponding author upon reasonable request.

